# A General Kinetic Model of Symporters Built on Cellular Homeostasis and Conformational Transitions

**DOI:** 10.64898/2026.10.15.759425

**Authors:** Wang Qiqian, Liang Feixue

**Affiliations:** School of Biomedical Engineering, Southern Medical University, Guangzhou, P.R. China

**Author notes:** Corresponding author: wang.

**Keywords:** symporter, mathematical modeling, conformational cycle, gating, probabilistic transition, kinetics, cellular homeostasis

## Abstract

General mathematical modeling of symport faces two fundamental challenges: (i) a unified kinetic description independent of protein-specific structural details; and (ii) quantitative cross-scale mapping between microscopic conformational transitions and macroscopic physiological observables. Grounded in the fundamental physiological principle of cellular homeostasis, we propose a unifying hypothesis for symporter gating: symporters actively regulate intracellular concentrations of specific solutes within thermodynamically feasible bounds. Based on this hypothesis, we construct a universal mathematical model of symporters, employing a minimal conformational cycle as the topological scaffold and probabilistic transition rates to quantitatively describe conformational interconversion kinetics. Model simulations of experimental data from the SGLT, NaPi-II, NKCC, and NBC transporter families yield consistent fits. The resulting parameters not only reproduce experimentally observed transport behaviors—but also reveal underlying mechanistic principles of symporter function.

## 1 Introduction

Secondary transporters are molecular machines that are ubiquitously present in every cell. Secondary transport is generally classified into symport and antiport. Symporters mediate symport—the transmembrane cotransport of substrates in the same direction.

Currently, in the theoretical modeling of transmembrane symport, although kinetic models have been developed for specific transport proteins, their applicability remains limited due to poor generalizability across different types of symporters. There is a pressing need for a general mathematical model—or equivalently, a universal mathematical framework—that enables quantitative characterization of symporter function. Such a framework should integrate multiple regulatory factors and be broadly applicable to diverse symport systems. It must not only capture the conformational state transition pathways but also elucidate the causal relationship between external signal inputs and the resulting transport behaviors. Consequently, the model should center on two key components: the regulatory mechanism governing the initiation and termination of transport, and the conformational dynamics underlying the symport process. Despite incomplete experimental data and structural information at present, existing knowledge is sufficiently robust to support the development of such a model.

To characterize the regulatory mechanism governing the initiation and termination of transport, it is essential to define: (i) the physiological conditions that activate symporters to initiate substrate translocation; and (ii) the conditions that trigger transport cessation or transition to a resting state.

The regulatory signals of symporters encompass a variety of types, such as ions, hormones, neurotransmitters, pH, and membrane potential. These signals can be categorized as either intracellular or extracellular based on their spatial localization. For example, during inward cotransport, a high intracellular concentration of substrates can trigger symporters to transition into a low-activity conformation, thereby preventing excessive substrate influx and preserving intracellular homeostasis. For SGLTs (sodium-glucose cotransporters), the rate-limiting step in forward transport under conditions of saturating ligand concentrations is the electroneutral dissociation of sodium ions from the cytosolic face of the transporter. In the case of SGLT1, the membrane potential, which exerts a directional effect on secondary active transport^1^, has a significant impact on the translocation of the empty carrier during the catalytic cycle, whereas its influence on other steps is relatively minor [2]. Additionally reversible inactivation induced by environmental factors, including extracellular pH, ion concentration, and temperature, can lead to conformational changes in transport proteins. Upon removal of these environmental stimuli, the protein conformation is restored, and functional activity is re-established.

On the other hand, systematic investigations into the conformational changes associated with the symport process have been conducted, resulting in a substantial body of detailed experimental data.

In recent years, the availability of high-resolution 2D and 3D structural data has significantly advanced our understanding of symporter molecular mechanisms. However, symport is an inherently dynamic process—one that cannot be fully captured or mechanistically deciphered from static structural snapshots alone. Consequently, despite the rapid accumulation of structural information, a general mathematical model capable of integrating this information to construct the dynamic process of symport remains elusive.

The alternating access model serves as a fundamental theoretical framework for understanding the mechanisms of secondary transport. Its conceptual foundation originates from early studies on membrane transport function [3]. Despite recent advancements in the field, this model remains scientifically valid. It is reinforced by the biochemical characteristics of transport reversibility and the underlying symmetry within all the MFS proteins [4].

This model categorizes the conformations of the transporter protein into two distinct types, T_e_ and T_i_: In T_e_, the polar cavity is open to the outer aqueous medium, but a part of the protein blocks access from the inner medium. In T_i_, the polar cavity is open to the inner aqueous medium, but a gate blocks access to the outer medium. [5] Evidently, this mechanism allows the transporter to block the ion transport, thereby enabling the selection of transmembrane ion flow direction.

During the transport process, upon sequential conformational changes in the carrier protein, the substrate binding sites are successively exposed to one side or the other of the membrane. This approach posits that the gate’s state fluctuates stochastically, with its opening and closing modeled as a Markov process [6]. This concept has since undergone further development and refinement through kinetic and biochemical studies, as well as theoretical analyses. These advancements have facilitated detailed descriptions of putative conformational events and related kinetic steps in specific carrier proteins. [7]

Previous mathematical models of symport, constrained by the absence of atomic structural data, failed to establish a direct connection with the molecular understanding of the symporter mechanism. Furthermore, the lack of essential components limited the models’ ability to effectively simulate the physiological experiments of symport. Fortunately, the increase in the number of available transporter protein structures in recent years has made our modeling work possible.

At present, it is widely accepted that symporters mediating substrate influx (referred to as *inward symporters* in this study) adopt at least three distinct conformational states: outward-open for substrate upload, substrate-bound occluded, and inward-open for substrate release. For Mhp1 and LeuT, all three conformations have been captured. Naturally, symporters that mediate substrate efflux (referred to as *outward symporters*) are also prevalent in biological systems. Accordingly, outward symporters should also exhibit at least three analogous conformational states: inward-open for substrate upload, substrate-bound occluded, and outward-open for substrate release. Therefore, to understand the transport mechanism, structural elucidation of a given transporter in at least three different conformations is required: the substrate-free outward-open state (Ce), the substrate-bound occluded state (C-Sc), and the substrate-free inward-open state (Ci). [8]

However, current understanding of dynamic changes in carrier proteins during the transport catalytic cycle remains limited. For instance, in the transport cycle, how many distinguishable conformational intermediates need to be considered, and how long do these intermediates remain within the cycle? In the carrier structures corresponding to the endpoints of these conformational changes, evidence of two conformational events was found, namely the opening or closing of the narrow gate, and the coordinated conformational transition between the Ce and Ci states. The conformational change that transforms a transporter from outward-to inward-facing is likely to go through one or more intermediate states. [9]

## 2 Foundation

Throughout this paper, vectors 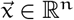 are by default *row vectors* (i.e., 1 × *n* matrices).

### Definition 1

**(Hadamard or Schur Product)** *For any two vectors* 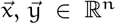, *their* component-wise product *(also known as Hadamard product or Schur product)* 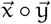 *is the vector in* ℝ^n^ *defined by* 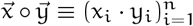.

### Definition 2

**(Hadamard Division)** *For any two vectors* 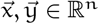, *their* component-wise division *(or Hadamard division)* 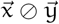 *is the vector in* ℝ^n^ *defined by* 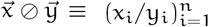.

### Definition 3

**(Hadamard Power)** *For any two vectors* 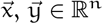, *their* component-wise power *(or Hadamard power)* 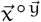 *is the vector in* ℝ^n^ *defined by* 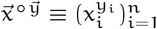.

### Definition 4

**(Broadcasted Power)** *For a scalar* x *∈* ℝ *and a vector* 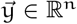, *the* broadcasted power *(or scalar-vector Hadamard power)* 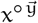 *is defined componentwise by* 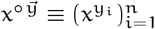.

### Definition 5

**(Component-wise Extension)** *For a function* f : *X → Y and arguments* x_1_, … , x_k_ *∈* X *(each a scalar or row vector), the* component-wise extension *of* f *in stacked notation applies* f *to each argument independently and stacks the results vertically:* f(*x*_1_; … ; *x*_k_) *≡* [f(*x*_1_); … ; f(*x*_k_)] . *When* f *is scalar-valued, this yields a column vector; when* f *is vector-valued, it yields a block column matrix*.

### Definition 6

**(Permutation)** *Two vectors* 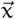 *and* 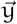 *are said to be* permutations *of each other if one can be obtained by rearranging the elements of the other. In particular*, 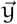 *is called a* permutation *of* 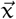, *denoted as* 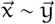.

Solute particles can be divided into two categories: charged and uncharged types. Among these, the charged particles are mainly ions. The movement of ions gives rise to an observable electric current. These charged species are capable of generating an electric field, which in turn influences ion migration and leads to an uneven distribution of intra- and extracellular ions. For example, in a hyperpolarized cell, positively charged ions inside the cell will move away from the membrane surface, while negatively charged ions will accumulate on the inner surface of the membrane. Numerous earlier models either neglected or inadequately addressed the issue of ion distribution, thereby limiting their overall accuracy and applicability.

Assume that the extracellular space is infinitely vast. Let the average intracellular concentration of solute particle ι be denoted by 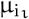 (conventionally written as [ι]_i_), the average extracellular concentration of ι by 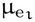 (written as [ι]_e_), and the density (per unit area) in the process of transmembrane transport of ι by s_mι_. For an ideal spherical cell with a radius of r_m_, assuming its cell membrane is homogeneous, the densities on the inner and outer surfaces of ι (written as s_iι_ and s_eι_) of the membrane can be estimated based on the average particle concentrations inside and outside the membrane:

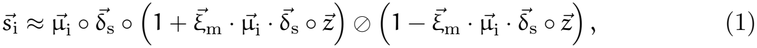

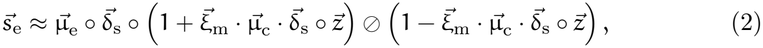

where δ_sι_ is the diameter of ι, z_ι_ is the valence of ι,

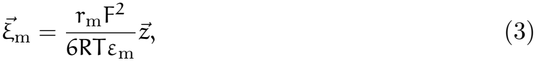

ε_m_ is the membrane permittivity and

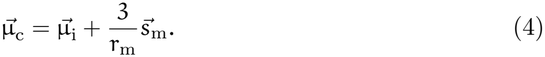

The derivation of Equations 1 to 4 [10] refers to the theory of the electrical double layer [11]. Considering a cell, we can estimate its membrane potential based on intracellular particle concentrations,

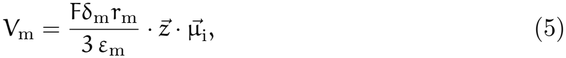

where δ_m_ is the membrane thickness.

Certain substances, particularly non-polar or weakly polar uncharged small molecules, can traverse the phospholipid bilayer of the cell membrane (denoted as *D*) directly without relying on membrane proteins. This mechanism is referred to as *simple diffusion*.

Molecules with high lipid solubility, such as benzene and steroids, exploit their hydrophobic nature to dissolve into the hydrophobic core formed by phospholipid tails and rapidly traverse the membrane. These molecules exhibit high affinity for the lipid bilayer and consequently diffuse at a relatively fast rate. Gaseous molecules such as oxygen and carbon dioxide, owing to their exceptionally high lipid solubility, readily dissolve in the lipid bilayer and diffuse rapidly. This process plays a critical role in respiratory gas exchange, particularly in the transfer of gases across the alveolar-capillary interface.

According to Fick’s law of diffusion, the diffusion flux 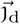—the amount of substance passing through a unit area of membrane per unit time—is directly proportional to the concentration difference across the membrane and is also influenced by the electric field force for charged species. Using the previously introduced estimation methods (see Equations 1 to 4), it follows that

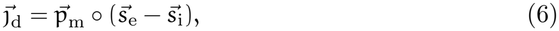

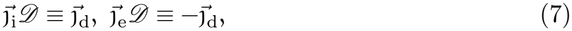

where the quantities 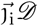 and 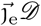 denote the material fluxes at the intracellular and extracellular surfaces, respectively, arising from simple diffusion, with the convention that flux is positive in the direction away from the membrane surface; 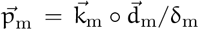 denotes the permeability coefficient, 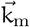 denotes the lipid-water partition coefficient, d_m_ is the diffusion coefficient, and δ_m_ represents the effective membrane thickness—an operational measure influenced by membrane fluidity rather than geometric dimensions alone. Thus, structural alterations in the membrane can significantly impair the efficiency of simple diffusion. In pathological conditions such as pulmonary fibrosis, the alveolar-capillary membrane thickens due to connective tissue hyperplasia and is frequently associated with reduced surfactant levels and decreased membrane fluidity, resulting in an increased effective thickness δ_m_ and a reduced diffusion coefficient 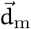, thereby diminishing the diffusion rates of oxygen and carbon dioxide.

## 3 Analysis

Conformation refers to the three-dimensional spatial arrangement of atoms within a protein molecule. The conformational changes of symporters enable the function of transporting substrates.

### Proposition 1

*A complete symport cycle can be decomposed into a series of strictly sequential steps. Each step has two endpoints, which are designated as stages in this study*.

The alternating access model serves as a fundamental theoretical framework for elucidating the mechanisms of secondary transport, with its conceptual foundation originating from early studies on membrane transport function [3]. Despite recent advancements in the field, this model remains scientifically valid and is reinforced by the biochemical characteristics of transport reversibility and the underlying symmetry within all the MFS proteins [4]. In this model, during the transport process, upon sequential conformational changes in the carrier protein, the substrate binding sites are successively exposed to one side or the other of the membrane, a concept that has since undergone further development and refinement through kinetic and biochemical studies as well as theoretical analyses, which have facilitated detailed descriptions of putative conformational events and related kinetic steps in specific carrier proteins [7].

The current understanding of the dynamic changes in carrier proteins during the transport catalytic cycle remains limited. For example, it is still unclear how many distinguishable conformational intermediates must be considered in the transport cycle and how long these intermediates persist within the process.

Conformational changes are continuous, but must be discretized in the mathematical model. As a result, not all intermediates can be considered: some are indistinguishable; others have change rates that cannot be reliably measured; and still others do not satisfy the significance criterion for inclusion. Nevertheless, certain conformational intermediates must be explicitly accounted for.

### Proposition 2

*The branch structures within the conformational transition network must be preserved*.

For instance, when two different substrates compete for the same binding site on a symporter protein, the system enters a critical branch point situated at the bifurcation of the conformational transition pathway.

This stage represents a dynamic equilibrium in which the proteins structural evolution proceeds along distinct trajectories depending on the identity of the bound substrate, thereby giving rise to divergent kinetic behaviors during subsequent transport steps. A well-documented example is the sodium/glucose cotransporter (S-GLT), in which the same binding site binds structurally similar yet functionally distinct substrates—such as glucose and galactose—eliciting different conformational rearrangements. These differences manifest as variations in transition rates, ion coupling efficiency, and overall transport kinetics, extending beyond mere quantitative disparities to potentially alter the functional outcome of the transport cycle. Omission of these essential stages—or some key intermediate states within the bifurcation process—from the mechanistic model would result in failure to account for the critical branching dynamics underlying substrate selectivity and transport efficiency. Hence, retaining such bifurcation structures is not merely an enhancement of model resolution; it is fundamental to ensuring predictive accuracy and biological fidelity in computational analyses of transporter dynamics.

### Proposition 3

*During symport, the carrier protein sequentially binds its substrates on one side of the membrane and translocates them across the membrane, releasing them on the opposite side, in a specific order*.

Based on a review of prior research [7, 12], a transport catalytic cycle can typically be categorized into the following stages:

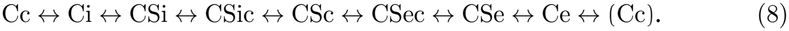

When the substrate and co-substrate are differentiated, the classification of stages becomes more complex:

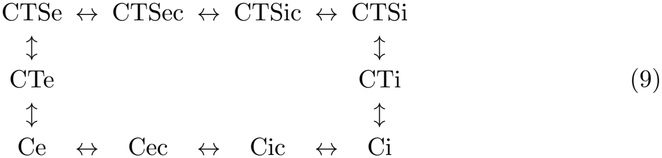

where C denotes the carrier protein, S denotes the substrate, and T denotes the cosubstrate; e and i correspond to the externally and internally exposed conformations of the carrier, respectively, while c indicates the occluded state. The binding order of the co-substrate and substrate follows a defined sequence, with T binding first, followed by S. This cycle consists of ten distinct stages (Cec, Ce, … , Ci, … , Cic), which are sequentially interconnected to form ten consecutive steps.

In simple terms, during the initiation phase of the transport cycle in inward (outward) symport, substrates bind to the carrier protein, which undergoes a conformational transition from the Ce (Ci) state to the CSec (CSic) state. Upon completion of the cycle, the carrier protein returns to its initial ground state through a substrate-free transition from the Ci (Ce) state back to the Ce (Ci) state, thereby enabling subsequent rounds of inward (outward) transport. [12]

That is to say, in the classical mechanism, during symport, the binding and release of substrate and co-substrate occur sequentially, such that the binding sequence is opposite to the dissociation sequence. Furthermore, transition CSic *↔* CSc *↔* CSec may occur stochastically with defined probabilities.

Nevertheless, certain studies suggest that the release order need not satisfy strict reversibility with respect to the binding order. For example, in hSGLT2-mediated inward cotransport, Na^+^ dissociation may precede 4FDG^2^ release [13]. However, this does not compromise the validity or construction of our mathematical model, which incorporates this factor.

### Proposition 4

*The gates of symporters are specific (sub)domains within transport proteins that, under defined conditions, regulate the accessibility of particular stages— a process referred to as gating*.

The concept of “gating” which refers to the opening or closing of a channel pore in response to a particular stimulus, was originally introduced in the field of ion channel modeling. It describes a gate within a channel as a structural component of the pore that prevents ion passage when the channel is in the closed or inactivated state and undergoes a conformational change upon stimulation, thereby allowing ions to traverse the channel. This terminology has since been extended to carrier proteins; however, due to functional differences between transporters and channels, the concept has generally been interpreted in two distinct ways: The first definition characterizes a “gate” in carrier function as a particular (sub)domain of the transporter that regulates the accessibility of the substrate binding site to its surroundings. The second definition broadens the scope of “gating” to include all conformational transitions of carrier protein binding and releasing substrate(s). [12]

Based on the integration of the aforementioned perspectives, it can be concluded that the gates of symporters are specialized (sub)domains within transport proteins that play a critical role in regulating the translocation of substrates across biological membranes. Given that the sequence of substrate binding and release is strictly conserved during the transport cycle, these gating elements—under specific biochemical or biophysical conditions, such as substrate binding, transmembrane ion gradients, or changes in membrane potential—function exclusively to permit or restrict access to key conformational intermediates. This precisely regulated process constitutes the mechanism known as gating.

### Proposition 5

*For any symporter, it can be posited that there are two distinct gates: one regulates the exposure of the binding site to a specific substrate, while the other governs the conformational transition of the substrate-bound protein between outward- and inward-facing states*.

In the carrier structures corresponding to the endpoints of the conformational changes, evidence of two conformational events was found, namely the opening or closing of the thin gate, and the coordinated conformational transition between the Ce and Ci states. The conformational change that transforms a transporter from outward-to inward-facing is likely to go through one or more intermediate states. [9] A thin gate refers to a part of the protein that regulates the accessibility of substrates. The opening or closing of thin gates is not correlated with significant changes in the whole protein. In contrast, a thick gate is typically assigned to a core component of the carrier structure that undergoes substantial conformational changes when the transporter switches between the Ce and Ci states. More specifically, some studies suggest that the thick gate’s opening and closing control the transition between CSec and CSic. [14, 12] In this context, the gates can be defined as structural barriers that mediate access between the substrate and the intracellular or extracellular environments, thereby preventing the establishment of dynamic equilibrium between bound and free substrate. The gating mechanism in symporters refers to the regulated control of these barriers.

### Proposition 6

*Each step in a transport cycle can be conceptualized as a chemical reaction. The transition rate of conformational state during this process is governed by the ion density at the membrane interface and the underlying gating mechanism*.

Any step can be decomposed into a combination of elementary steps, which are categorized into three types: the substrate binding step, analogous to a bimolecular binding reaction; the pure conformational change step, analogous to a process with a constant transition rate; and the substrate release step, analogous to a molecular dissociation reaction. Notably, conformational changes occur not only during pure conformational transitions but also during both substrate binding and release processes. The transition rate is governed by the ion density on the inner and outer surfaces of the membrane, provided that the protein is accessible to the substrate in both the initial and final states.

### Proposition 7

*Symporters regulate intracellular concentrations of specific substances via transmembrane transport to maintain cellular homeostasis*.

Cell homeostasis is a crucial concept in biology and medicine, defined as the ability of cells to maintain a stable internal environment. [15] Central to this ability is the regulation of intracellular substance concentrations (especially the ion concentration), which ensures physiological stability through coordinated control of transmembrane influx and efflux—a condition primarily attributable to the functional activity of transport proteins.

In prior research, we developed a general mathematical model for ion channels [10]. This model posits that the primary function of ion channel activity is to maintain specific intracellular ion concentrations, and it has demonstrated satisfactory performance in various physiological simulation experiments [10, 16]. Given that the stability of intracellular solution concentrations arises from the combined effects of both active and passive transport mechanisms, we have reason to believe that the evolutionary origin of symporters was primarily aimed at preserving specific intracellular substance concentrations. As will be demonstrated in the subsequent discussion, this fundamental objective alone is capable of giving rise to a wide array of complex physiological phenomena.

### Proposition 8

*If an action induces a deviation of the intracellular concentration of a specific solute from its steady-state value and is sensed by a symporter, it constitutes a physiological stimulus for that symporter. In response to such stimuli, symporters mediate solute transport—specifically inward cotransport or outward cotransport—to preserve cellular homeostasis*.

Stimulation denotes a physiological or environmental perturbation imposed on a cell that induces a measurable change in its biochemical state—especially its intra-cellular fluid—thereby disrupting its homeostatic equilibrium.

For a perturbation to elicit a directed transport response, the resulting deviation in intracellular solute concentration must be sensed by the transport machinery and transduced into a measurable change in translocation flux.

Evidence from several symporter families indicates that ligand- or state-dependent sensing occurs, although the molecular mechanisms underlying it differ in how directly the intracellular state is coupled to the transport cycle. Voltage- and charge-coupled sensing constitutes a comparatively direct mode of signal transduction [17]. In contrast, upstream signaling cascades constitute an indirect mode, where sensing and execution can be modularly separated: the deviation is sensed by a dedicated sensor and relayed to the symporter by post-translational modification ^3^. In such functionally coupled configurations, a symporter and its upstream signal-sensing components—including specific receptors, sensor kinases, and ligand-binding domains— constitute an integrated molecular functional unit, designated as a *symporter system*.

The aforementioned signals are fundamentally determined by the intracellular distribution of solutes. Given that symporters respond exclusively to stimuli acting at the membrane interface, the relevant stimulus is more precisely defined by the local concentration of specific solutes at the intracellular face of the membrane.

## 4 Model

### 4.1 Conformation

Based on Proposition 1, during the transport process, the state of symporter *C* — responsible for providing binding sites and herein defined as a *stage* (denoted by ψ)—undergoes dynamic transitions.

#### Definition 7

*A transition from one stage* ψ *to another* ψ^*′*^ *is defined as a* directed step (ψ, ψ^*′*^).

#### Definition 8

*Each step* {ψ, ψ^*′*^} *is bidirectional, encompassing a pair of mutually inverse directed steps* (ψ, ψ^*′*^) *and* (ψ^*′*^, ψ).

In the special case where ψ = ψ^*′*^, the step constitutes a self-loop. A complete cycle (denoted as ϕ) consists of a sequential series of such directed steps. Within each cycle, the binding sites are alternately exposed to either side of the membrane (see Proposition 3), a mechanism referred to as alternating access. Therefore, every conformation can be uniquely mapped to a corresponding (ψ, ϕ) pair. When the analysis is confined to substrate transport and other functional roles of the protein are excluded, the conformation can be adequately represented by (ψ, ϕ).

The function of the carrier protein is mediated by transitions among multiple conformational states. Consequently, a symporter *C* may be formally defined as the ensemble of all conformations it can adopt under physiological conditions. In this framework, the complete set of all conformations of a symporter constitutes the protein itself. Thus, when (ψ, ϕ) represents a valid conformational state of *C* , we express this relationship by stating that (ψ, ϕ) *∈ C*.

Let the set of all symporters on a membrane *M* that are in the conformational state (ψ, ϕ) be denoted by *ℳ* {(ψ, ϕ)}, which satisfies the distributive property with respect to union,

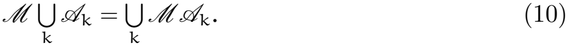

Thus, the set of symporters *C* on membrane *M* is given by *ℳ C* . Let |*ℳ* | denote the area of *ℳ* , and let

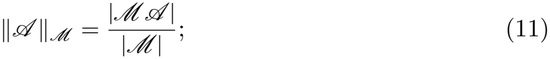

then the density of *ℳ* {(ψ, ϕ)} is given by

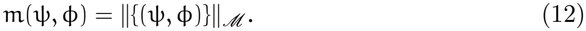

Thus, the density of symporters *C* on membrane *ℳ* is *∥C ∥*_*ℳ*_ , abbreviated as *∥C ∥* for brevity. There holds

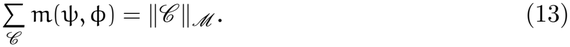

Furthermore, the transition flow from (ψ, ϕ) to (ψ^*′*^, ϕ) is expressed as

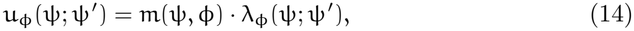

and the transition flow from (ψ, ϕ) to (ψ, ϕ^*′*^) is expressed as

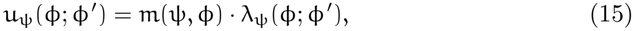

where λ_ϕ_(ψ; ψ^*′*^) denotes the transition rate from (ψ, ϕ) to (ψ^*′*^, ϕ), and λ_ψ_(ϕ; ϕ^*′*^) denotes the transition rate from (ψ, ϕ) to (ψ, ϕ^*′*^). Let

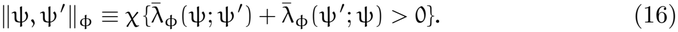

If *∥*ψ, ψ^*′*^*∥*_ϕ_ = 1, then it follows that there exists a transition step {ψ, ψ^*′*^} under the condition ϕ; otherwise, if *∥*ψ, ψ^*′*^*∥*_ϕ_ = 0, no such step exists. The transition flow can be employed to calculate the temporal changes in the state density of symporters,

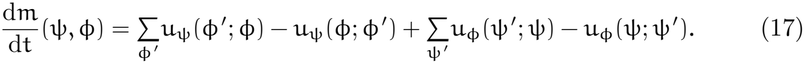

### 4.2 Cycle

It is well established that within a single transport cycle, a symporter may cotransport multiple substrates.^4^ Building upon Equations 8 and 9, we propose a more general carrier model for the symport mechanism:

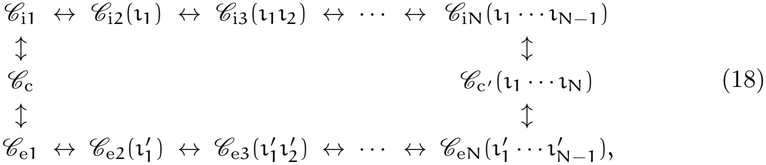

where *C*_ii_ and *C*_ei_ represent the inward-facing and outward-facing conformations of carrier *C* that expose the i-th binding site to the intracellular and extracellular sides, respectively; *C*_c_ represents a fully occluded, substrate-free intermediate state between *C*_i1_ and 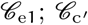 represents a fully occluded, substrate-bound intermediate state between *C*_iN_ and *C*_eN_; N represents the total number of binding sites in *C* ; and ι_n_ and 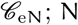 represent the substrates bound to the n-th inward- and outward-facing binding sites, respectively, satisfying 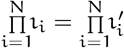.

We refer to Equation 18 as a complete transport cycle. It should be noted that certain transport proteins can participate in multiple distinct transport cycles. From Proposition 6, the entire cycle can be represented by a set of corresponding chemical equations:

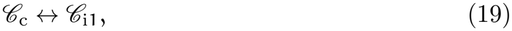

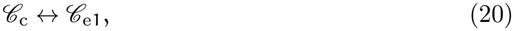

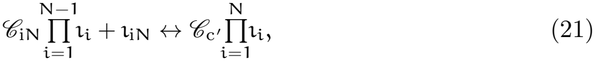

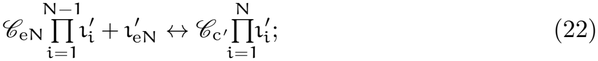

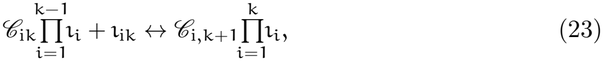

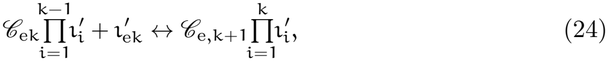

for k = 1, *· · ·* , N − 1; where 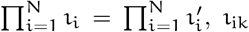 represents the intracellular 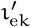 represents the extracellular 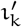, and the empty-product convention is adopted: 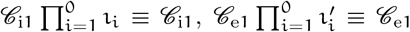. The standard format of the aforementioned Equations 19 to 24 can be formally expressed as follows:

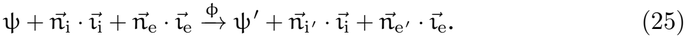

Here, ψ and ψ^*′*^ represent the stages (associated with bound substrate), with the constraint that a unique transition path exists from ψ to ψ^*′*^. The variables 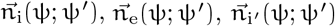 and 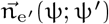 represent the ratios of the respective substances involved in the conformational transition, while 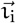 and 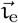 correspond to the vectors describing the solute particle compositions inside and outside the cell, respectively. Building upon the general form of the rate equation and incorporating the Hill equation [19], we formulate the following expression to estimate the transition rate from (ψ, ϕ) to (ψ^*′*^, ϕ), denoted as λ_ϕ_(ψ, ψ^*′*^), which is given by

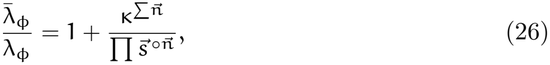

where 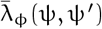 represents the maximum achievable transition rate; κ(ψ, ψ^*′*^) is the equidensity half-saturation constant from (ψ, ϕ) to (ψ^*′*^, ϕ) for all 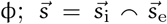 represents the concatenated vector of intracellular and extracellular particle concentrations; and 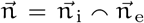. Then, the particle fluxes across the intracellular and extracellular membrane surfaces mediated by *C* = *p × f* are

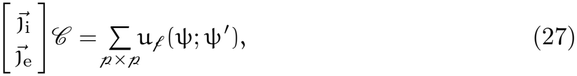

where

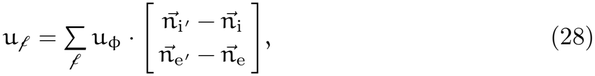

and where negative values correspond to substrate entry into the symporter, while positive values correspond to substrate exit from the symporter. Thus,

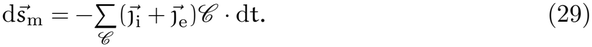

The transport rate of *C* can be obtained from

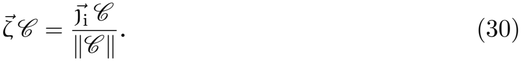

### 4.3 Gating

The gating mechanism regulates the accessibility of specific stages (see Proposition 4), where 0 and 1 represent inaccessibility and accessibility, respectively. Specifically, this mechanism governs the parameter λ_ϕ_. Consequently, a gating function g_ϕ_(ψ; ψ^*′*^) *∈* {0, 1} can be defined such that

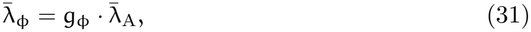

where 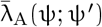 is the transition rate from ψ to ψ^*′*^ when ψ^*′*^ is accessible to the substrate(s).

According to Proposition 7, the symporter (system) must exhibit selectivity for the target particle(s). We posit that this selectivity arises from structurally defined sensitive site(s). For carrier-intrinsic sensing, these sites are embedded in *C* ; for cases in which sensing and execution are modularly separated, they reside in the sensor component of an upstream sensor–transducer apparatus. Given the particle density on the inner membrane surface, 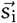 (see Equation 1), particles adsorb onto the sensitive site(s) of the symporter system *C* with a finite probability. Let the adsorption equilibrium constant for the specific particle ι at the sensitive site(s) of *C* be æ_*C* ι_. The sensitive site(s) of *C* do not form long-lived bound states; consequently, the system operates in the low-coverage regime far from monolayer saturation, where the fraction of free sites is approximately unity. Thus, the total adsorbed particle density on the sensitive site(s) satisfies

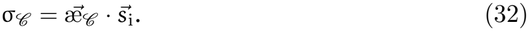

We assume that *C* functions to maintain σ_*C*_ at 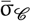. As stated in Proposition 8, the condition 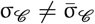 constitutes a stimulus on *C* . The intensity of this stimulus is quantified by the magnitude of deviation between σ_*C*_ and 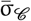, and is formally defined as 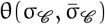. The stimulus intensity θ should exhibit scale covariance with respect to the particle density σ, i.e., there exists a function f such that

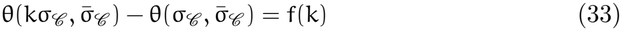

for all k > 0. Additionally, θ satisfies the reflexivity condition θ(x, x) = 0 for all x > 0. If θ is continuously differentiable on (0,) *×* (0,), then it admits the logarithmic form

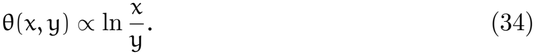

It is known that conformational gating dynamically modulates the active site accessibility, thereby permitting or restricting substrate binding (see Proposition 5). From Proposition 8, substrate binding is inhibited when 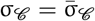, corresponding to the *unactivated* state of the carrier. Let the unactivated state be denoted by a = 0; the corresponding transport cycle is then expressed as ϕ|_a=0_. Under this condition, *C*_c_ is a sink vertex in the cycle; that is,

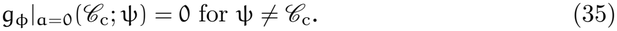

When 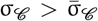, the symporters tend to promote the *outward cotransport*, i.e., the efflux of specific particles. Such symporters are designated as *outward-permissive* and are formally denoted by a = 1. In contrast, when 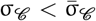, the symporters tend to promote the *inward cotransport*, i.e., the influx of specific particles. Such symporters are designated as *inward-permissive* and are formally denoted by a = −1.

Let 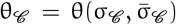. Because the binding site of an unactivated *C* is neither accessible to any substrate nor occupied by one, it follows necessarily that ψ = *C*_c_. Therefore, for θ_*C*_ > 0, the carrier in state (*C*_c_, ϕ|_a=0_) undergoes a transition to (*C*_c_, ϕ|_a=1_); for θ_*C*_ < 0, the carrier in state (*C*_c_, ϕ|_a=0_) undergoes a transition to (*C*_c_, ϕ|_a=−1_). Conversely, when θ_*C*_ = 0, the carrier in state (*C*_c_, ϕ|_a=*±*1_) transitions to (*C*_c_, ϕ|_a=0_).

The transition rate from (*C*_c_, ϕ|_a=0_) to (*C*_c_, ϕ|_a=*±*1_) defines the activation rate of *C* . The activation rate depends on stimulus intensity; we therefore denote it by

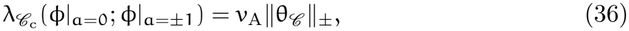

where the domain of ν_A_ is the half-open interval [0, ∞). The Taylor series expansion of v_A_(x) about x = 0 yields

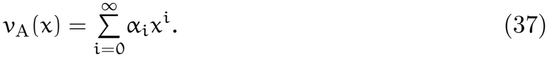

Since ν (0) = 0, it follows that α_0_ = 0. If 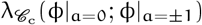 is smooth at θ = 0, then α_1_ = 0. Thus, in the vicinity of 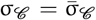, the relationship admits a quadratic polynomial approximation, given by

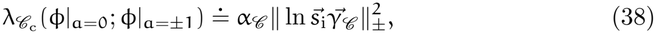

where α_*C*_ denotes the *activation coefficient* of *C* , and 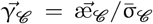 denotes the *sensitivity coefficient* of *C*.

This transition is reversible. Activated symporters exhibit an inherent tendency to return to their unactivated state, ensuring that upon removal of the stimulus, they revert to the initial state, corresponding to the resting state of the transport proteins when the cell is in homeostasis. Within our modeling framework, the process is more rigorously formulated as a transition from (*C*_c_, ϕ|_a=*±*1_) to (*C*_c_, ϕ|_a=0_). Should this transition rate remain constant throughout (denoted as ν_−A*C*_), it follows that

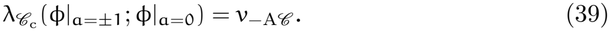

From Propositions 5 and 6, a gating mechanism is required to inhibit transport mediated by symporters with a = 1 during inward transport or by symporters with a = −1 during outward transport. Moreover, such interruption must occur precisely at the step between the outward- and inward-facing states—namely, at 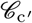. Let ϕ|_b=1_ denote the inhibited transport, and let ϕ|_b=0_ denote the uninhibited transport. Under this condition, *C*_c_ ^*′*^ is a source vertex in the cycle; that is,

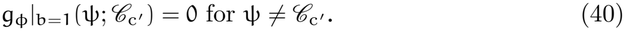

In what follows, denote ϕ(x, y) *≡* ϕ|_a=x,b=y_. Except for the cases referenced in Equations 35 and 40, g_ϕ_(ψ, ψ^*′*^) = 1 holds universally. Consequently, we have

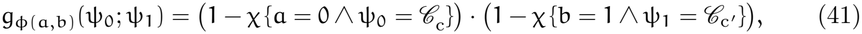

where *χ* is the indicator function. It follows that 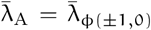 and Equation 31 admits a more precise expression:

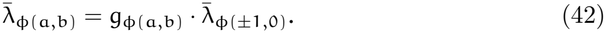

Given that a symporter cannot simultaneously undergo both inward and outward transport, the transition from *C*_eN_ to 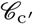 corresponds to inward transport, whereas the transition from *C*_iN_ to 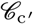 corresponds to outward transport. It follows that

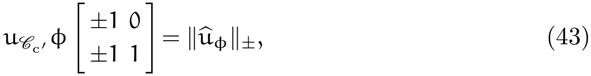

where

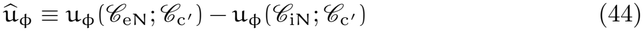

is the net transition flow at 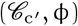. Any symporter *C* that has been interrupted and has released all bound substrates must undergo the state transition from b = 1 to b = 0, with the rate of this process given by v_−B*C*_ , after returning to the state a = 0, before it can fulfill the necessary conditions to complete the subsequent transport cycle. This relationship is formally described by the following equation:

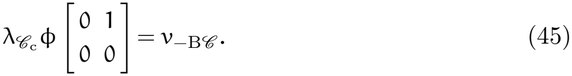

### 4.4 Simplification

In practice, it is difficult to accurately determine the transition parameter 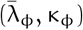 for each step {ψ, ψ^*′*^}. When the division of steps is too fine, a step may occur on an extremely short timescale or exhibit significant duration variability, which arises from stochasticity. Therefore, in model calculations, multiple steps—particularly those with extremely fast rates—can be concatenated without compromising the accuracy of the computation or the validity of electrophysiological experiment simulations.

#### Definition 9

*Given two distinct steps s*_1_ ≠ *s*_2_, *if s*_1_ *∩ s*_2_ *∅, then s*_1_ *and s*_2_ *are said to be adjacent, and any element* ψ *∈ s*_1_ *∩ s*_2_ *is referred to as an intermediate state between s*_1_ *and s*_2_.

Given steps {ψ_1_, ψ_2_} and {ψ_2_, ψ_3_}, since {ψ_1_, ψ_2_} *∩* {ψ_2_, ψ_3_} = {ψ_2_}, it follows that {ψ_1_, ψ_2_} and {ψ_2_, ψ_3_} are adjacent, and ψ_2_ is their intermediate state.

#### Definition 10

*The operation defined by s*_1_ *⊕ s*_2_ *≡ s*_1_ *∪ s*_2_ − *s*_1_ *∩ s*_2_ *is designated as the merging of s*_1_ *and s*_2_.

The merging of steps {ψ_1_, ψ_2_} and {ψ_2_, ψ_3_} is {ψ_1_, ψ_3_}. This operation entails the omission of the intermediate stage ψ_2_, combining two adjacent steps into a single, unified transition.

This approach is employed to simplify state transition sequences and eliminate redundant intermediate states. By removing the shared intermediate node ψ_2_, a more streamlined state transition path is obtained, thereby reducing the representational complexity of the system and enhancing analytical efficiency. The resulting step emphasizes the direct relationship between the initial and final states while attenuating the significance of intermediate transitional stages, making it suitable for applications where fine-grained internal dynamics are not the primary focus.

In certain cases, transfer cycles are sufficiently complex that the merging of all adjacent steps is not feasible; equivalently, not all intermediate states may be omitted.

#### Definition 11

*Given* ϕ, *The degree of* ψ *is defined as* deg 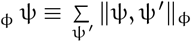.

#### Definition 12

*If* deg_ϕ_ ψ > 2, *then* ψ *is defined as a branch point on* ϕ, *and the set* {*s* : ϕ *∈ s*} *constitutes a branch passing through* ϕ.

According to Proposition 2, given ϕ, if ψ is a branch point on ϕ, then for the reduced 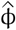 , it necessarily follows that 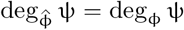 This implies that all stages associated with such branch points are indispensable. Moreover, even if a given stage does not itself correspond to a branch point, it remains essential whenever its removal undermines the structural integrity of an adjacent branch—specifically, by collapsing two or more otherwise distinct conformational pathways into a single trajectory. Furthermore, since the ψ values of inactive *C* are uniformly equal to *C*_c_, the stages satisfying ψ = *C*_c_ are consistently preserved.

Let there be a sequence of steps, *s*_i_ = {ψ_i−1_, ψ_i_} for i = 1 to n. Since *s*_k_ and *s*_k+1_ are adjacent with the intermediate transitional stage given by ψ_k_ for all k < n, successively merging them yields

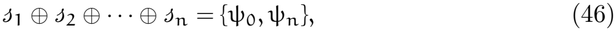

which is denoted as 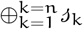 hereinafter. For {ψ_0_, ψ_n_}, according to Equation 25, the reaction equation is given by

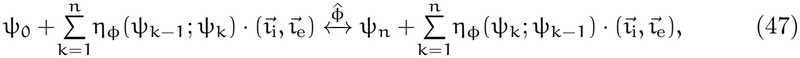

where 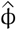 denotes the simplified transfer cycle obtained through the integration of the steps from *s*_1_ to *s*_n_. The transition rate of the merged 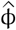 can still be estimated using Equation 27, where 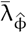 represents a newly calibrated empirical parameter. The steps omitted in 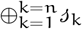 are not required to be computed within 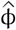; thus, we set

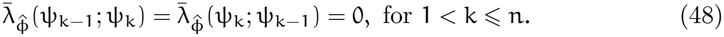

However, these steps can be regarded as approximately exhibiting uniform flow behavior, implying that 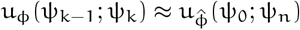 for k = 1 to *n*.

In Equation 44, the net transition flow at 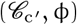 is computed as 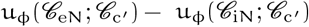. When deg 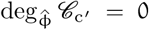, there exist integers k, k^*′*^ *∈* {1, 2, … , N} such that 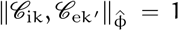. Under this condition, the approximations 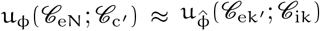 and 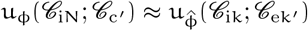 hold, yielding an analogous estimate for the net transition flow:

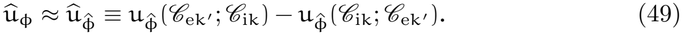

When 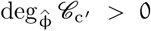 but 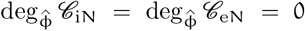, it follows that there exist k, k^*′*^ *∈* {1, 2, … , N − 1} such that 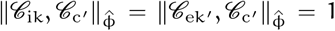. In this case, we have 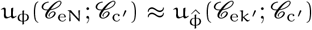 and 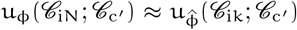, which yields an analogous estimation formula for the net transition flow:

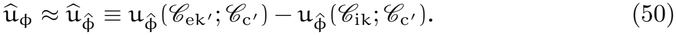

### 4.5 Constraints

Thermodynamic constraints are universal and fundamental: their applicability is independent of the structural features of specific transport proteins, species origin, or cell type, and arises solely from the laws of energy conservation and entropy increase. All processes involving transmembrane transport of substances—including all symport mechanisms—are necessarily subject to thermodynamic constraints.

In this study, the net free energy change associated with symport is determined by the difference in electrochemical potential across the membrane, which dictates transport directionality. For any *C* = *p × f*, symport mediated by *C* necessarily satisfies thermodynamic constraints for all ϕ *∈ f*. Therefore, it is assumed here that *C* always remains in the activated state (a ≠ 0 and b = 0), allowing the carrier to continuously mediate the symport cycle. When a system is isolated, thermo-dynamic constraints compel it to evolve toward equilibrium. Therefore, sustained carrier-mediated transport drives the cellular state—comprising intracellular and extracellular solute concentrations as well as the conformational distribution of the carrier—toward equilibrium.

Thus, if all other transmembrane transport mechanisms are excluded and the constraints 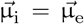 and 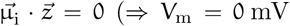 and 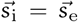, i.e. 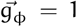 are imposed, the conformational distribution of *C* converges to a steady state and the net transition flow vanishes. This steady-state distribution constitutes a thermodynamic equilibrium of the system. At this balance limit, for any simple transport cycle of *C*

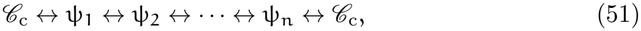

the following detailed-balance condition must hold:

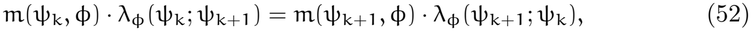

for k = 0, 1, … , n, where ψ_0_ = ψ_n+1_ = *C*_c_. It follows that

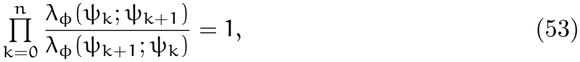

under the thermodynamic constraints 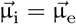 and 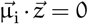. Taking the limit µ ⟶ ∞, where 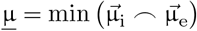, we obtain

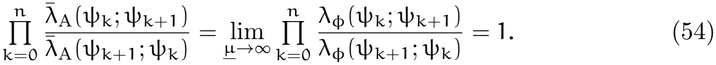

Since Equation 53 holds for all 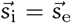, and Equation 54 establishes the asymptotic invariance of the cycle product, it follows that

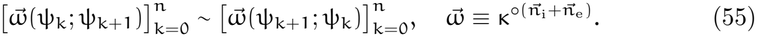

The constraints expressed in Equations 54 and 55 impose thermodynamically necessary restrictions on the model parameters, thereby guaranteeing that, at thermodynamic equilibrium, the system exhibits both time-invariant macroscopic properties and the absence of any macroscopic material circulation.

### 4.6 Boundaries

When the experiment *X* is formally initiated at time t = t_0_, it is reasonable to assume that the external solution concentration remains constant and unperturbed for all t < t_0_,

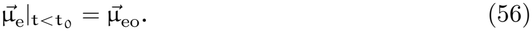

This assumption establishes a well-defined initial condition, which is essential for constructing mathematical models and analyzing the dynamics of transmembrane transport. Moreover, the specification of the intracellular solution as t *→* − ∞ is required,

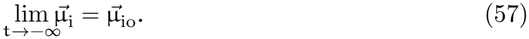

Under this condition, the system has sufficient time to reach a steady state or quasi-equilibrium, in which the concentrations of various species, electrochemical gradients, and the distribution of transporter protein states remain invariant over time. Specifically, we may further assume that at t *→* − ∞, all symporters *C* on the membrane *M* reside in their initial states, as

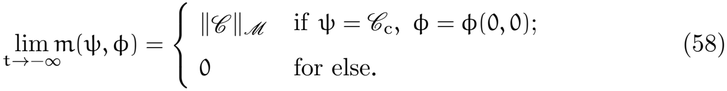

The experiment formally begins at t ⩾ t_0_. During this phase, it may be necessary to actively modulate the concentrations of intracellular or extracellular solutions in order to simulate specific physiological or pathological conditions, or to investigate the response mechanisms of transmembrane transport processes to external stimuli.

This study does not account for the narrow intercellular gaps and assumes that the extracellular space is sufficiently extensive to effectively buffer local concentration fluctuations. Under this assumption, changes in extracellular solution concentration are primarily governed by the external perfusion system, rather than by cellular metabolic activities or diffusive interactions between neighboring cells. This idealization simplifies the model’s boundary conditions and is justified under most in vitro experimental settings ^5^. The perfusion rate is denoted as 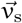, and the concentration of the perfusion solution is 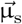. Under the assumptions of this paper, in regions far from the outer surface of the membrane where the electric field vanishes, the perfusion-induced change in extracellular concentration can be calculated as

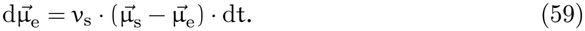

In particular, if v_s_ *→* ∞, then 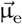 reaches 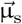 instantaneously. In the following calculations, unless otherwise specified by experimental constraints, it is assumed that v_s_ is sufficiently large; this assumption will not be restated.

For the regulation of intracellular fluid, patch-clamp technology is a widely used and highly significant experimental technique. In patch-clamp experiments, particle flow arises primarily through two physical mechanisms: concentration-driven diffusion and electromigration. To simplify the theoretical model and emphasize dominant factors, it is commonly assumed that the volume of the pipette is extremely limited— such as at the tip of a microelectrode or within the localized region in tight contact with the cell membrane—where ionic concentration gradients rapidly homogenize. Under this condition, the contribution of concentration diffusion to the total ion current becomes negligible and can therefore be neglected. This simplification not only reduces the complexity of mathematical modeling but also facilitates a clearer understanding of electric field-driven ion transport. In this regime, the dominant mechanism governing particle flow is electromigration—the directional movement of charged ions under an externally applied electric field. This study focuses on silver electrodes as a representative example. Silver electrodes are extensively employed in electrophysiological measurements owing to their excellent conductivity, stable electrochemical performance, and ease of fabrication. In particular, when configured as Ag/AgCl electrodes, the redox reaction on the surface is

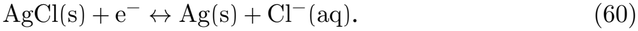

Thus, we derive the differential change in intracellular solute concentration as

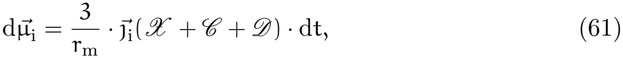

where

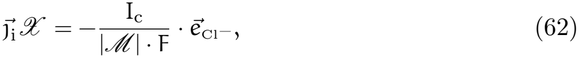

I_c_ denotes the electrode current. The relationship between the electrode current and the clamping potential is given by

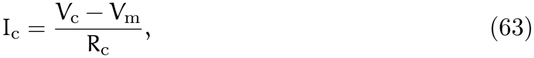

where R_c_ represents the resistance of the pipette when filled with solution.

The aforementioned method is equally applicable to other types of metal electrodes. Despite variations in electrochemical properties among these materials, they share a consistent methodology with respect to the fundamental ion current detection mechanism and the mode of electrical connection with cells.

## 5 Results

This study employs the aforementioned mathematical model to simulate canonical physiological experiments. These simulations serve to demonstrate the model’s practical applicability and to validate its theoretical and empirical soundness. All simulation data and results are presented in the article, including figures, the main text, and supplementary notes. No new physiological experimental data were generated in this study.

### 5.1 Sodium-glucose symport

SGLTs mediate glucose uptake against its concentration gradient by coupling glucose transport to the downhill movement of sodium ions along their electrochemical gradient. This mechanism was first proposed in 1960 [20] and represents a landmark discovery in membrane transport biology. Subsequently, the gene encoding SGLT1 was cloned—a pivotal advance in molecular physiology [21]. Further studies have identified multiple SGLT isoforms, including SGLT2 and SGLT3.^6^ Given SGLT1’s paradigmatic role in elucidating Na^+^-coupled symport mechanisms, this study applies the mathematical model developed herein to simulate key physiological functions of the transporter, which serves as the first case of numerical experiments in this paper.

In each transport cycle of SGLT1, two sodium ions and one glucose molecule are stoichiometrically coupled and translocated in the same direction across the membrane. Substrate binding occurs in a strictly ordered, sequential manner—Na^+^ binds first, followed by glucose, which serves as the final ligand to complete the ternary complex [22]. Therefore, SGLT1-mediated transport activity can be monitored as a sodium current generated by glucose, and the kinetic model parameters can be determined by recording the currents as a function of membrane potential and intracellular and extracellular concentrations of glucose and sodium ions.

In the present numerical simulation, the transport cycle of SGLT1 ^7^ can be decomposed into steps (*C* = SGLT1): ^8^

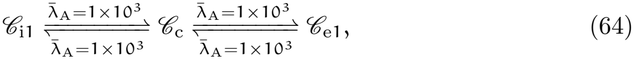

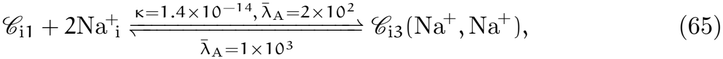

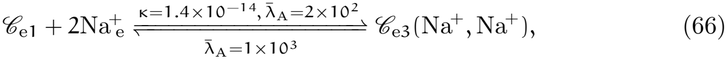

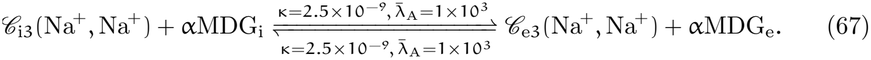

In standard electrophysiological assays using *Xenopus* oocytes heterologously expressing human SGLT1 (hSGLT1), the membrane potential is initially clamped at −50 mV to establish a defined baseline for studying Na^+^-coupled inward glucose transport. Subsequently, the membrane potential is stepped to a series of test voltages across a physiologically and experimentally feasible range, and steady-state transporter-associated currents are measured to construct voltage–current relationships. Systematic variation of extracellular Na^+^ and αMDG concentrations generates families of voltage–current relationships, permitting quantitative characterization of hSGLT1’s substrate affinity, voltage sensitivity, and coupled transport mechanism. Due to constraints imposed by oocyte membrane integrity and viability, the experimentally accessible voltage range is typically bounded between −150 mV and 50 mV. Physiological experiments demonstrate that, across a range of αMDG concentrations, hSGLT1-mediated voltage–current relationships exhibit robust voltage dependence: net inward current declines toward zero at depolarized potentials V_m_ > 50 mV, where-as at strongly hyperpolarized potentials V_m_ < 150 mV, current saturates (simulation results are presented in Figure 1). Notably, under hyperpolarized conditions (i.e., V_m_ = −150 mV), the maximal sugar transport rate mediated by hSGLT1 is independent of [Na^+^]_e_ (simulation results are presented in Figure 2B), whereas the maximal Na^+^ transport rate increases with rising [αMDG]_e_ (simulation results are presented in Figure 2A), indicating that glucose binding governs the rate-limiting step of the transport cycle when the electrical driving force is saturating. [23]

**Figure 1:**
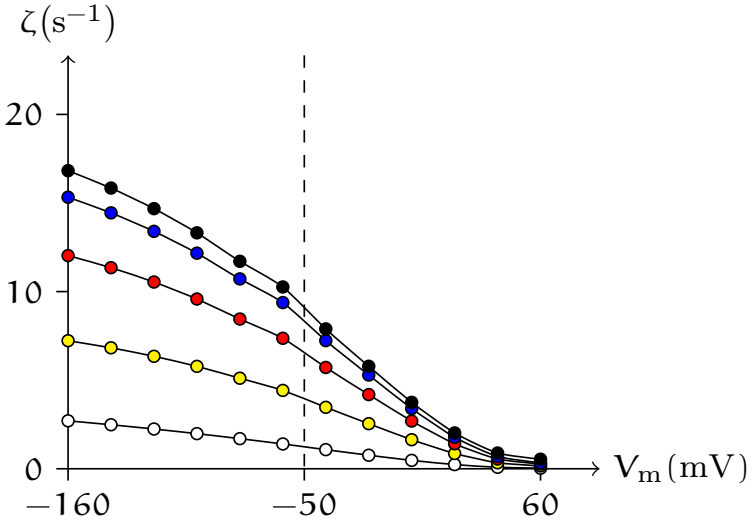
Computational simulation of the membrane potential-dependent glucose inward transport rate mediated by hSGLT1. Transport rate, 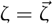 hSGLT1_αMDG_, with the inward direction defined as positive. The data of the white, yellow, red, blue, and black circles correspond to [αMDG]_e_ = 0.1, 0.7, 2, 5, and 10 mM respectively.

**Figure 2:**
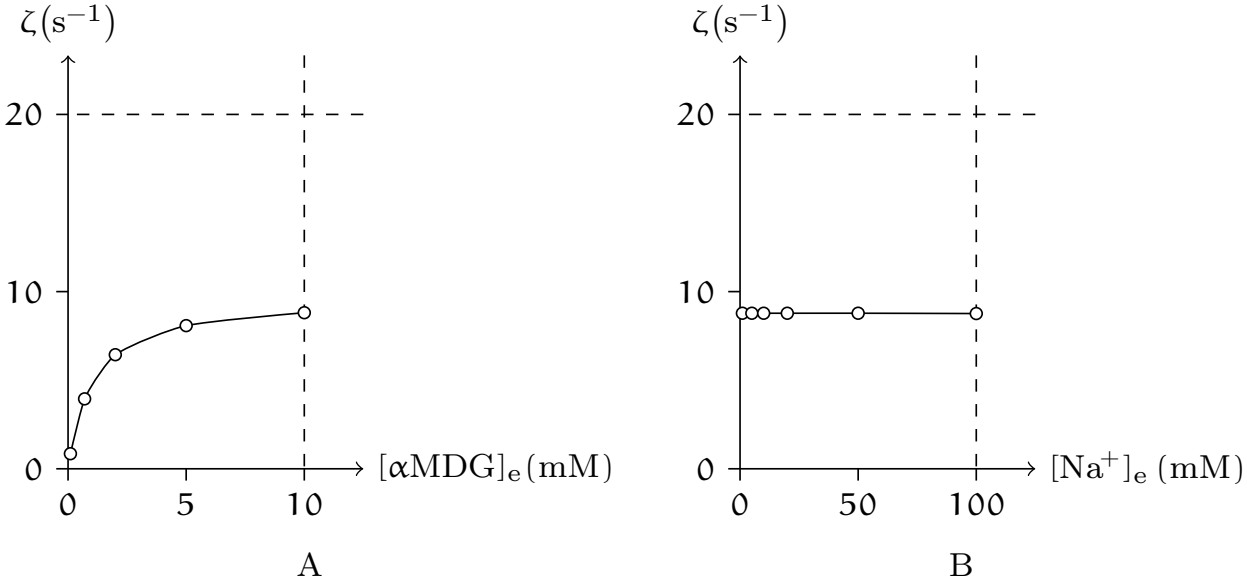
Simulated relationship between the extracellular solution concentration and the hSGLT1-mediated inward glucose transport rate. Transport rate, 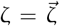 hSGLT1[αMDG], with the inward direction defined as positive. (A) Concentration-response relationship at V_m_ = −50 mV, corresponding to the experimental condition in Figure 1. (B) [Na^+^]_e_ was varied stepwise (1, 5, 10, 20, 50, 100 mM) while [αMDG]_e_ = 10 mM and V_m_ = −50 mV were held constant.

On the other hand, when intracellular sodium and sugar concentrations are high and extracellular concentrations are low (typically by setting [Na^+^]_e_ = 0 mM or [αMDG]_e_ = 0 mM), SGLT1 mediates outward cotransport, generating an outward current. In oocytes expressing rabbit SGLT1 (rSGLT1), holding the initial membrane potential at 0 mV while applying different cytoplasmic concentrations of αMDG yields a set of outward currents attributable to rSGLT1-mediated transport that increase with [αMDG]_i_ and saturate at 80 mM [24] (simulation results are presented in Figure 3).

**Figure 3:**
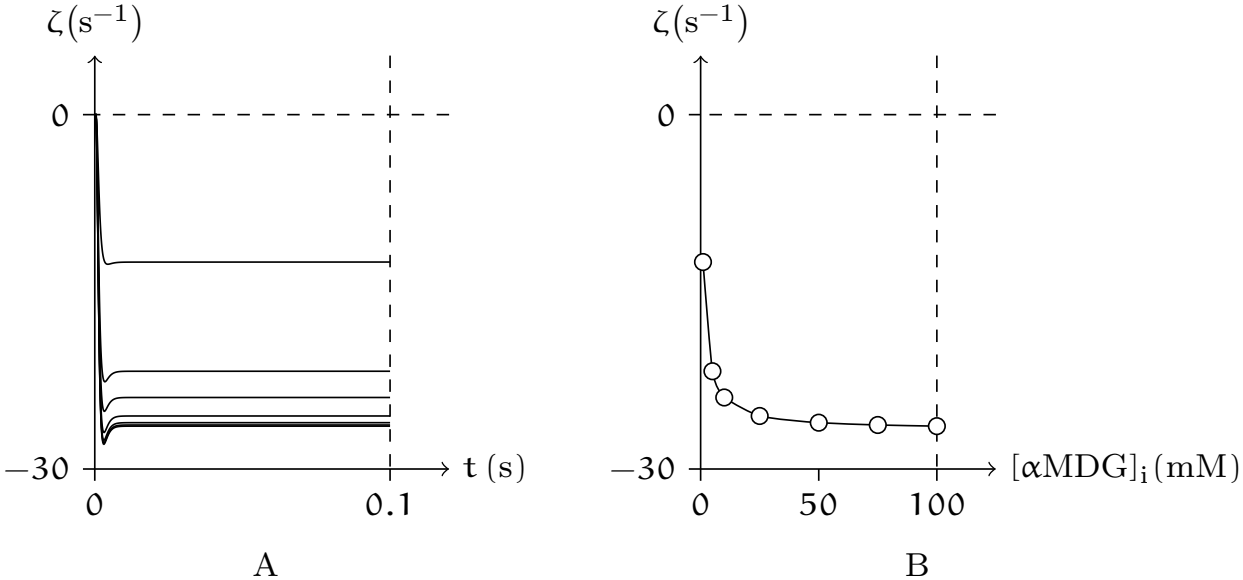
Numerical simulation of hSGLT1-mediated outward transport as a function of intracellular αMDG concentration under high intracellular sodium conditions. Transport rate, 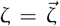 hSGLT1_αMDG_, with the inward direction defined as positive. (A) [Na^+^]_i_ = 100 mM; [αMDG]_i_ = 1, 5, 10, 25, 50, 75, 100 mM, with each concentration assigned sequentially to the transport curves in panel A, ordered from top to bottom. (B) Data were sampled at t = 0.1 s from the time course shown in panel A.

Next, we present a simulation of an illustrative example featuring a branch point (see Definition 12).

### 5.2 Sodium-coupled phosphate symport

NaPi-IIa and NaPi-IIb are electrogenic Na^+^-P_i_ symporters. In contrast, NaPi-IIc functions as an electroneutral symporter with a 2 : 1 (Na^+^ : P_i_) stoichiometry. This differs from the well-characterized 3 : 1 stoichiometry of NaPi-IIa/b. [25]

However, experimental evidence demonstrates that Li^+^ can functionally substitute for Na^+^ as the driving ion in NaPi-IIa/b. This substitution markedly perturbs the kinetics of downstream conformational transitions. A carrier model proposes a sequential symport mechanism: Li^+^ and Na^+^ compete for a putative high-affinity cation-binding site; occupancy by either ion permits subsequent cooperative binding of one Na^+^, one P_i_, and a third Na^+^—thereby completing the fully loaded transport complex [26]. According to this symport mechanism, and using NaPi-IIb as a representative example, our model yields numerical simulation results that are consistent with experimentally observed physiological data.

In the present numerical simulation, the transport cycle of NaPi-IIb ^9^ can be decomposed into steps (*C* = NaPi2b):

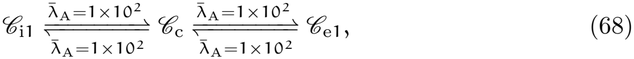

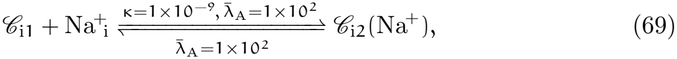

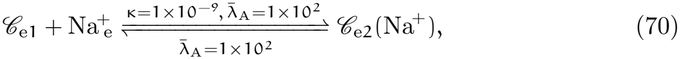

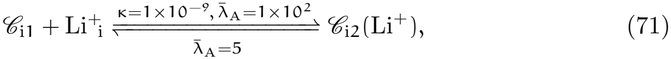

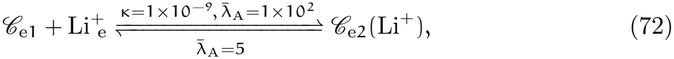

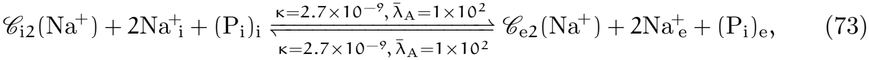

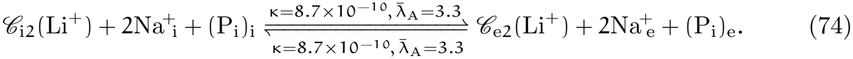

For oocytes expressing the NaPi-IIb isoforms, substituting 100 mM Na^+^ with 100 mM Li^+^ (marked as 100Li) in the incubation medium resulted in minimal phosphate uptake—approximately 3.0% of that measured in 100 mM Na^+^ (marked as 100Na). Replacing 50% of external NaCl with either LiCl (50 mM Na^+^, 50 mM Li^+^, marked as 50Na50Li) or choline Cl (50 mM Na^+^, 50 mM choline, marked as 50Na50Ch) revealed that Li^+^ reduced uptake by ∼ 50% relative to the 50Na50Ch control, indicating inhibition of the symporter.

To gain a more comprehensive understanding of NaPi-IIb function, the driving force and transport current can be measured (since NaPi-IIb is an electrogenic Na^+^-P_i_ symporter) under external solutions of varying monovalent cation concentration, with [P_i_]_e_ fixed at 1 mM. By clamping the membrane potential across a range of values, for each external solution, a membrane potential-transport current curve was obtained [27], where the transport current is proportional to the transport rate (simulation results are presented in Figure 4). For a given membrane potential *V*_m_, the dependence of transport rate on extracellular solution concentration was thus determined (simulation results are presented in Figure 5). Electrophysiological experiments showed that, at any membrane potential, when [Na^+^]_e_ > 25 mM, equivalent Li^+^ substitution further reduced the transport rate relative to the choline control. Moreover, the two conditions yielded markedly distinct response profiles. Under choline replacement, the transport rate increased with rising extracellular [Na] and gradually approached saturation (simulation results are presented in Figure 5D). In contrast, equimolar Li^+^ replacement produced a qualitatively different response: for [Na^+^]_e_ < 75 mM, the transport rate rises shallowly with [Na^+^]_e_; for [Na^+^]_e_ 75 mM, it increases approximately exponentially (simulation results are presented in Figure 5E). [26] This phenomenon occurs because, when [Li^+^]_e_ > 25 mM, most of the symporter *C* = NaPi2b is stabilized in step {*C*_i2_(Li^+^), *C*_e2_(Li^+^)}, whereas this population declines sharply as [Li^+^]_e_ decreases toward 0 mM (see Figure 6).

**Figure 4:**
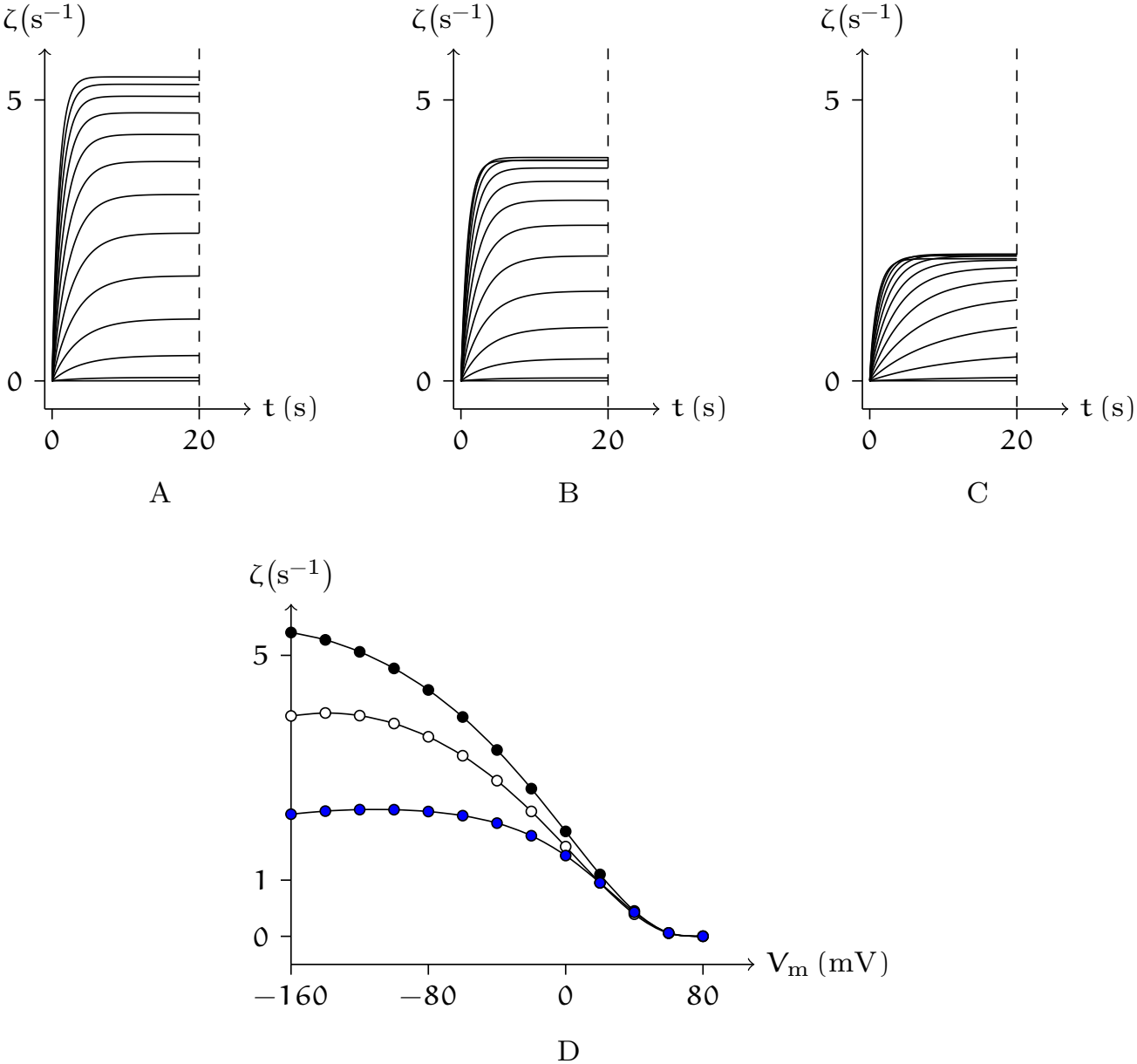
Computational simulation of the membrane potential-dependent phosphate transport rate mediated by NaPi-IIb. Transport rate, 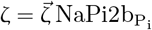, with the inward direction defined as positive. (A) Time-dependent change in NaPi2b-mediated phosphate transport rate upon activation in a 100Na extracellular solution. (B) Time-dependent change in NaPi2b-mediated phosphate transport rate upon activation in a 50Na50Ch extracellular solution. (C) Time-dependent change in NaPi2b-mediated phosphate transport rate upon activation in a 50Na50Li extracellular solution. In each panel, the superimposed traces— ordered from top to bottom—correspond to membrane potentials clamped at −160 mV to 80 mV in 20 mV increments. (D) Steady-state transport rates, quantified at t = 20 s post-activation in panels A, B, and C, are indicated by black, white, and blue circles, respectively representing the 100Na, 50Na50Ch, and 50Na50Li experimental conditions.

**Figure 5:**
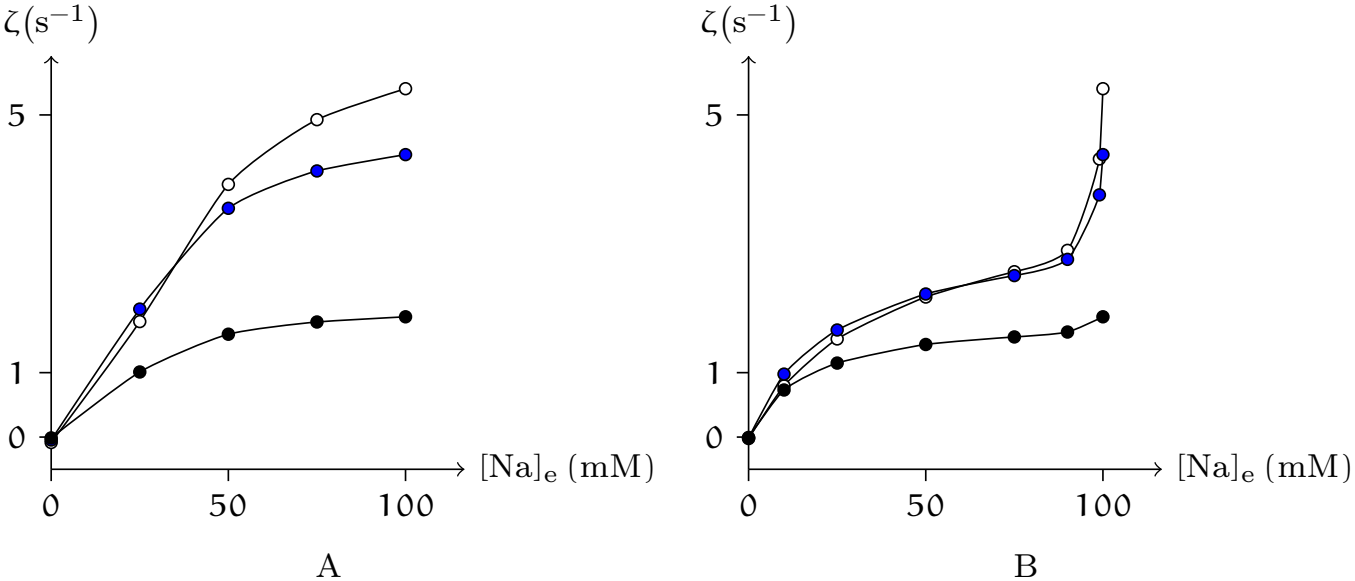
Numerical simulation of NaPi-IIb transport rates at different extracellular solution concentrations. Transport rate, 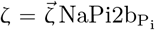, with the inward direction defined as positive. (A) [Na^+^]_e_ + [choline]_e_ = 100 mM. (B) [Na^+^]_e_ + [Li^+^]_e_ = 100 mM.

**Figure 6:**
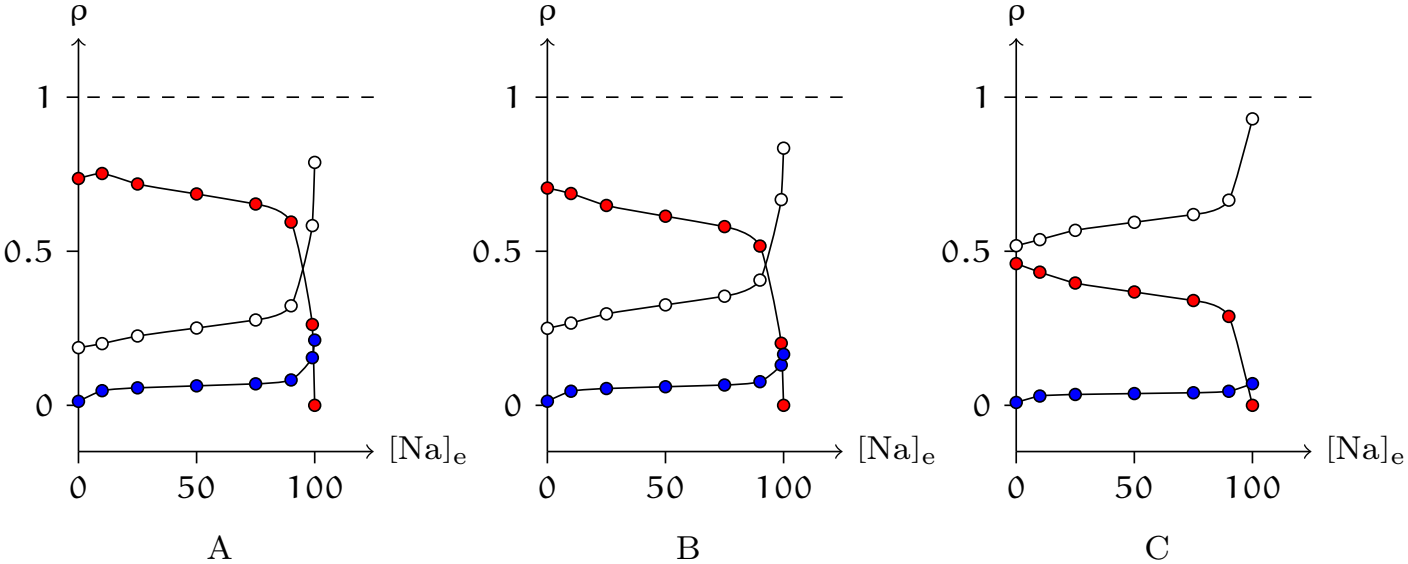
Effect of extracellular Li^+^ concentration on NaPi-IIb conformational dynamics: computational simulations under fixed total extracellular monovalent cation concentration ([Na^+^]_e_ + [Li^+^]_e_ = 100 mM). The red, blue, and white circles denote the respective proportions of {*C*_i2_(Li^+^), *C*_e2_(Li^+^)}, {*C*_i2_(Na^+^), *C*_e2_(Na^+^)}, and {*C*_i1_, *C*_e1_}, where *C* = NaPi2b. (A) Clamping voltage V_c_ = −160 mV. (B) V_c_ = −80 mV. (C) V_c_ = 0 mV.

In both aforementioned cases, the transport proteins mediate coupled electrogenic symport of exactly two substrate classes. To test the generalizability of our model, we now examine a representative electroneutral symporter capable of concurrently handling three substrate classes.

### 5.3 Sodium-potassium-chloride cotransport

Since the initial identification of the electroneutral Na^+^-K^+^-2Cl^−^ cotransport mechanism, this transport mode has been experimentally verified across diverse cell types. Two isoforms of the Na^+^-K^+^-2Cl^−^ cotransporter (NKCC)—NKCC1 and NKCC2— have been molecularly cloned and functionally characterized. NKCC1 exhibits broad cellular expression, having been detected in virtually all mammalian cell lineages examined thus far.

NKCC is generally regarded as an inward symporter that, under physiological conditions, mediates the electroneutral influx of Na^+^, K^+^, and 2Cl^−^ into the cell, thereby establishing and maintaining an elevated intracellular Cl^−^ concentration.

In the present numerical simulation, the transport cycle of NKCC ^10^ can be decomposed into steps (*C* = NKCC):

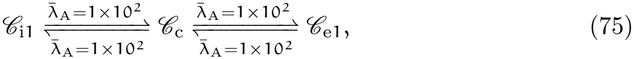

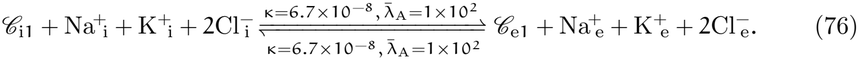

To investigate its inward transport properties, elevated extracellular concentrations of Na^+^, K^+^, and Cl^−^ are typically employed, such as [Na^+^]_e_ = 325 mM, [K^+^]_e_ = 100 mM, [Cl^−^]_e_ = 561 mM, [Mg^2+^]_e_ = 65 mM, and [Ca^2+^]_e_ = 3 mM; the intracellular environment is rendered sodium-free, for instance, by setting [K^+^]_i_ = 400 mM and [Cl^−^]_i_ + [glutamate^−^]_i_ = 400 mM. Varying [Cl^−^]_i_ allows for assessment of NKCC transport activity dependence on intracellular chloride concentration. The maximum inward chloride flux occurs when the intracellular chloride concentration [Cl^−^]_i_ is 0 mM, and the chloride influx mediated by NKCC decreases as [Cl^−^]_i_ increases. [28] The numerical simulation results of this behavior are presented in Figure 7.

**Figure 7:**
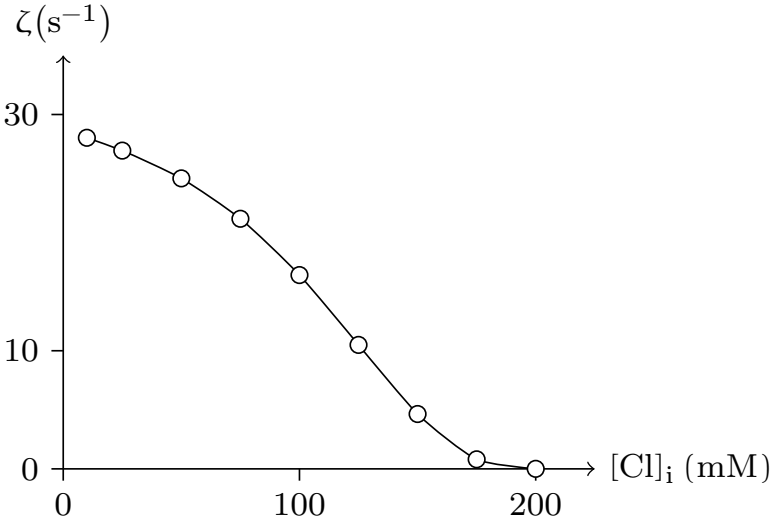
Numerical simulation of the steady-state transport activity of NKCC as a function of intracellular chloride concentration. Transport rate, 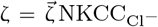, with the inward direction defined as positive.

To probe the ion-specific determinants of NKCC-mediated cotransport influx, [Cl^−^]_i_ = 0 mM is routinely established. This chloride-free intracellular condition permits unambiguous attribution of observed flux changes to variations in extracellular Na^+^, K^+^, and Cl^−^ concentrations (simulation results are presented in Figure 8). [29] While NKCC is predominantly characterized as an inwardly directed cotransporter, experimental conditions featuring high intracellular chloride (e.g., [Cl^−^]_i_ = 300 mM) coupled with low extracellular Na^+^ or Cl^−^ can elicit measurable outward NKCC-mediated flux [30, 31] (simulation results are presented in Figure 9).

**Figure 8:**
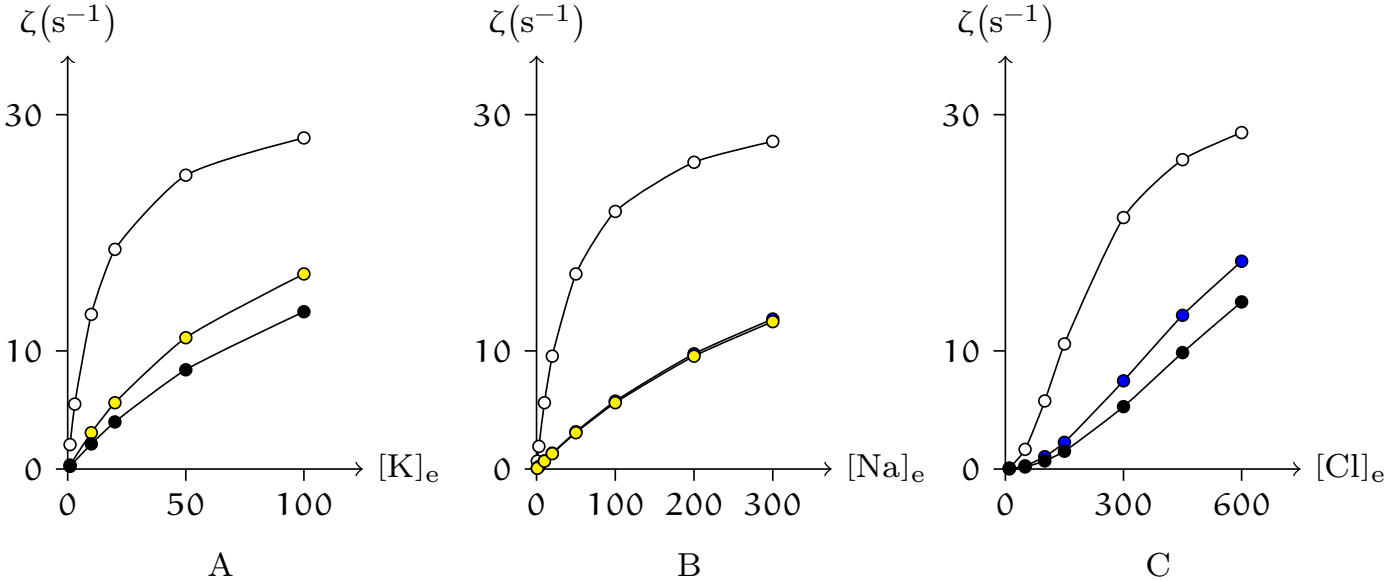
Numerical simulation of the steady-state transport activity of NKCC as a function of extracellular ionic concentration (mM). Transport rate, 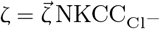, with the inward direction defined as positive. (A) White circles: [Na^+^]_e_ = 325 mM, [Cl^−^]_e_ = 561 mM; Yellow circles: [Na^+^]_e_ = 50 mM, [Cl^−^]_e_ = 561 mM; Black circles: [Na^+^]_e_ = 325 mM, [Cl^−^]_e_ = 180 mM. (B) White circles: [K^+^]_e_ = 100 mM, [Cl^−^]_e_ = 561 mM; Yellow circles: [K^+^]_e_ = 10 mM, [Cl^−^]_e_ = 561 mM; Blue circles: [K^+^]_e_ = 100 mM, [Cl^−^]_e_ = 180 mM. (C) White circles: [K^+^]_e_ = 100 mM, [Na^+^]_e_ = 325 mM; Blue circles: [K^+^]_e_ = 100 mM, [Na^+^]_e_ = 50 mM. Black circles: [K^+^]_e_ = 10 mM, [Na^+^]_e_ = 325 mM.

**Figure 9:**
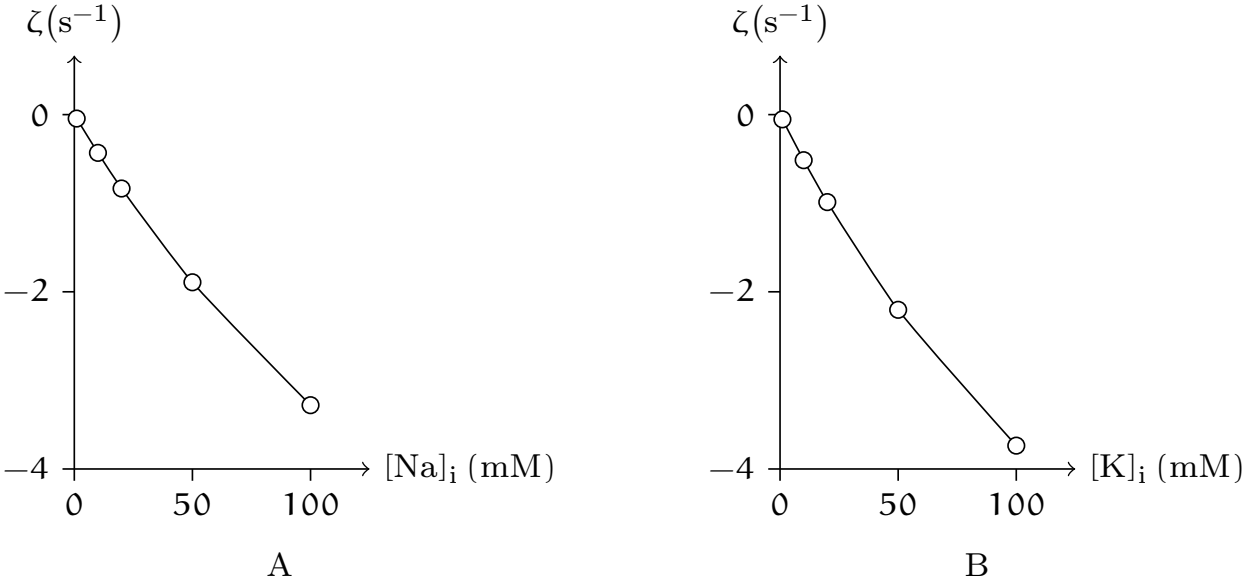
Numerical simulation of NKCC-mediated ion efflux. (A) [K^+^]_i_ = 50 mM, [Cl^−^]_i_ = 300 mM. (B) [Na^+^]_i_ = 60 mM, [Cl^−^]_i_ = 300 mM. Transport rate, 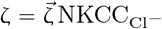, with the inward direction defined as positive.

Because almost every cellular process is sensitive to changes in intracellular pH, the regulation of pH_i_ is essential for maintaining normal cellular function. Below, we apply our model to compute a case in which pH_i_ is a relevant factor.

### 5.4 Sodium-bicarbonate cotransport

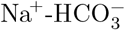cotransporters (NBC) function as either acid extruders or acid loaders. Below, we simulate a physiological experiment in which Rat kidney NBC (rkNBC) is heterologously expressed in *Xenopus* oocytes. NBC stoichiometry in *Xenopus* oocytes is 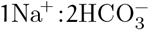, implicating a possible accessory protein interaction [32].

In the present numerical simulation, the transport cycle of rkNBC ^11^ can be decomposed into steps (*C* = rkNBC):

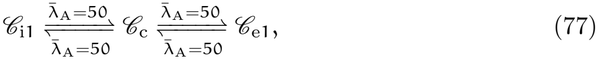

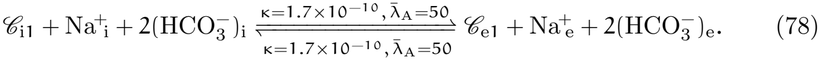

Functional properties of rkNBC expressed in *Xenopus* oocytes are as follows: rkN-BC expression attenuates 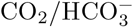-induced intracellular acidification and accelerates pH_i_ recovery. Upon 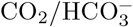 application, rkNBC-expressing oocytes exhibit rapid hyperpolarization, whereas control oocytes undergo slow depolarizationboth responses being attributable to electrogenic 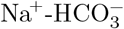 influx or its absence. Removal of extracellular Na^+^ induces intracellular acidification and large depolarization in rkNBC oocytes, consistent with reversal of inward cotransport (Na^+^-coupled HCO^−^ efflux). All observed effects are fully reversible upon Na^+^ restoration, confirming that rkNBC function strictly requires extracellular Na^+^. [33, 32]

The numerical simulations of the above phenomena and their functional characteristics are presented in Figure 10. These properties exhibit highly coupled, nonlinear dynamic behavior whose temporal evolution is governed by the interplay of intra-cellular pH buffering capacity, membrane capacitance, transmembrane ion gradients, and rkNBC’s intrinsic transport kinetics. Consequently, mechanistic attribution to any single molecular determinant or biophysical parameter remains unreliable when based solely on qualitative observation. However, our mathematical model explicitly incorporates the transport rate (see Figure 11A) and gating variables of rkNBC (see Figure 11B–D), and concurrently solves for the time-dependent behavior of other ionic currents (see Figure 12)—based on the established framework of passive ion transmembrane transport [10]—thereby enabling dissection of the causal relationships underlying the observed electrophysiological responses.

**Figure 10:**
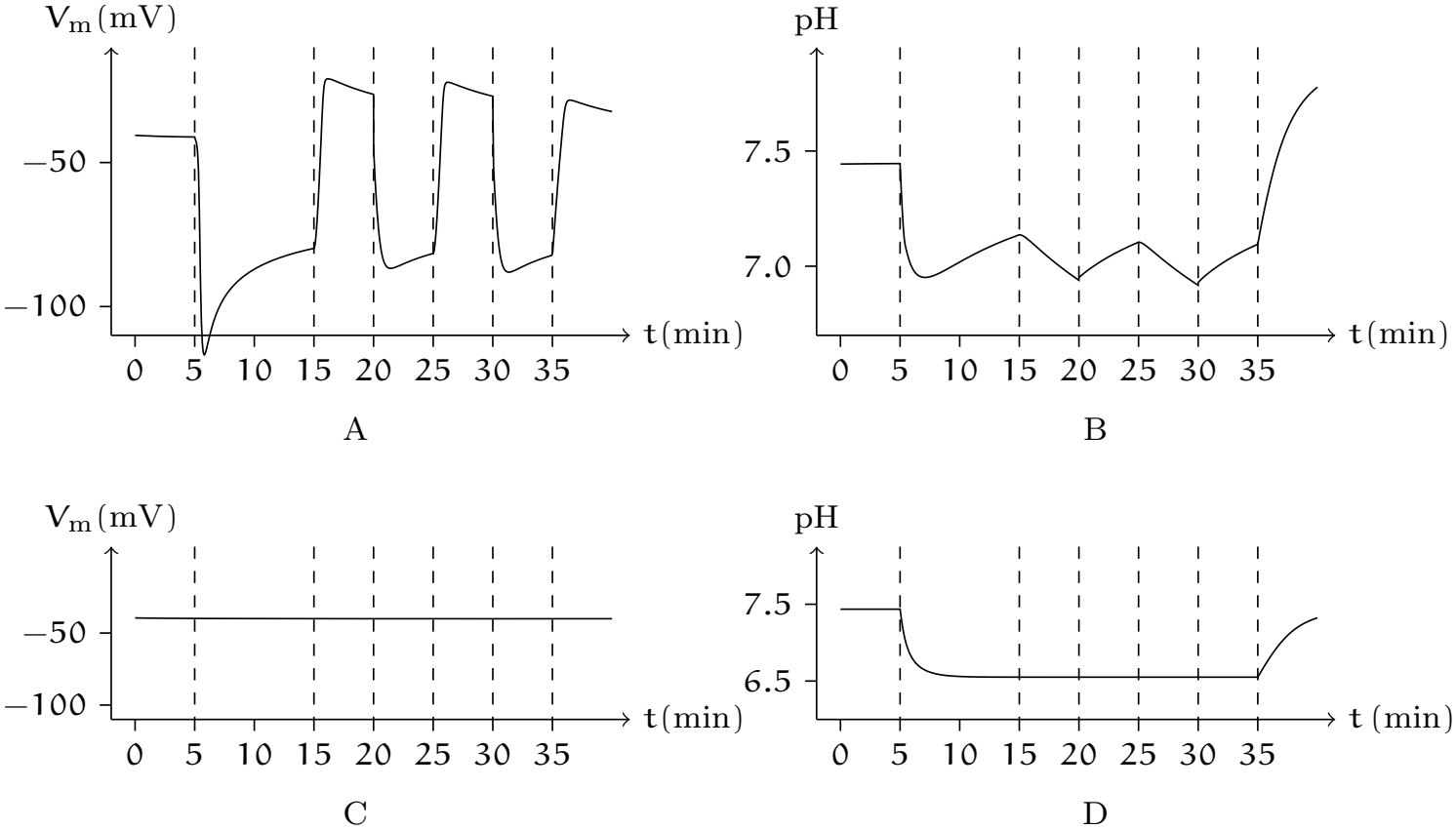
Numerical simulation of the physiological experiment in which rkNBC is heterologously expressed in *Xenopus* oocytes. The initial intracellular fluid consisted of 80 mM KCl and 10 mM NaCl, with pH adjusted to approximately 7.4. The initial extracellular fluid consisted of 100 mM NaCl, with pH adjusted to approximately 7.4. For t/(60s) in [5, 35], 1.5% CO_2_/10 mM 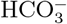 was added to the extracellular fluid. For t/(60s) in [15, 20] *∪* [25, 30], Na^+^ was removed from the extracellular fluid. Here, V_m_ denotes the membrane potential, and pH denotes the intracellular pH value. Numerical simulations of oocyte expression of rkNBC (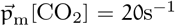, ∥rkNBC∥ = 2 nmol/m^2^) are shown in panels A and B. Numerical simulations of control oocyte (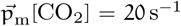, ∥rkNBC∥ = 0 nmol/m^2^) are shown in panels C and D.

**Figure 11:**
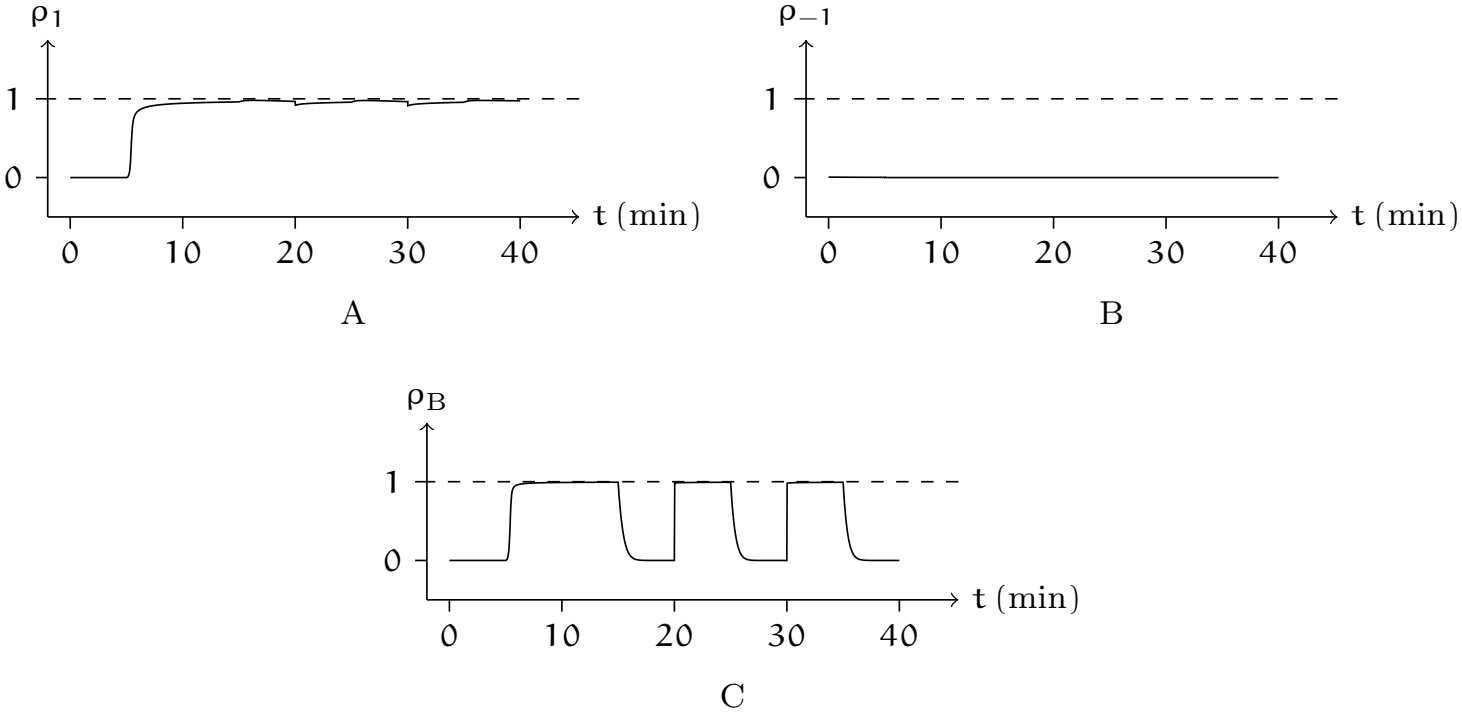
The transport rate and gating variables of rkNBC, as explicitly defined and dynamically updated within the numerical simulation. The data in this figure and in panels A and B of Figure 10 were obtained from the same numerical simulation. (A) Fraction of the outward-permissive state; (B) Fraction of the inward-permissive state; (C) Fraction of the interrupted transport state.

**Figure 12:**
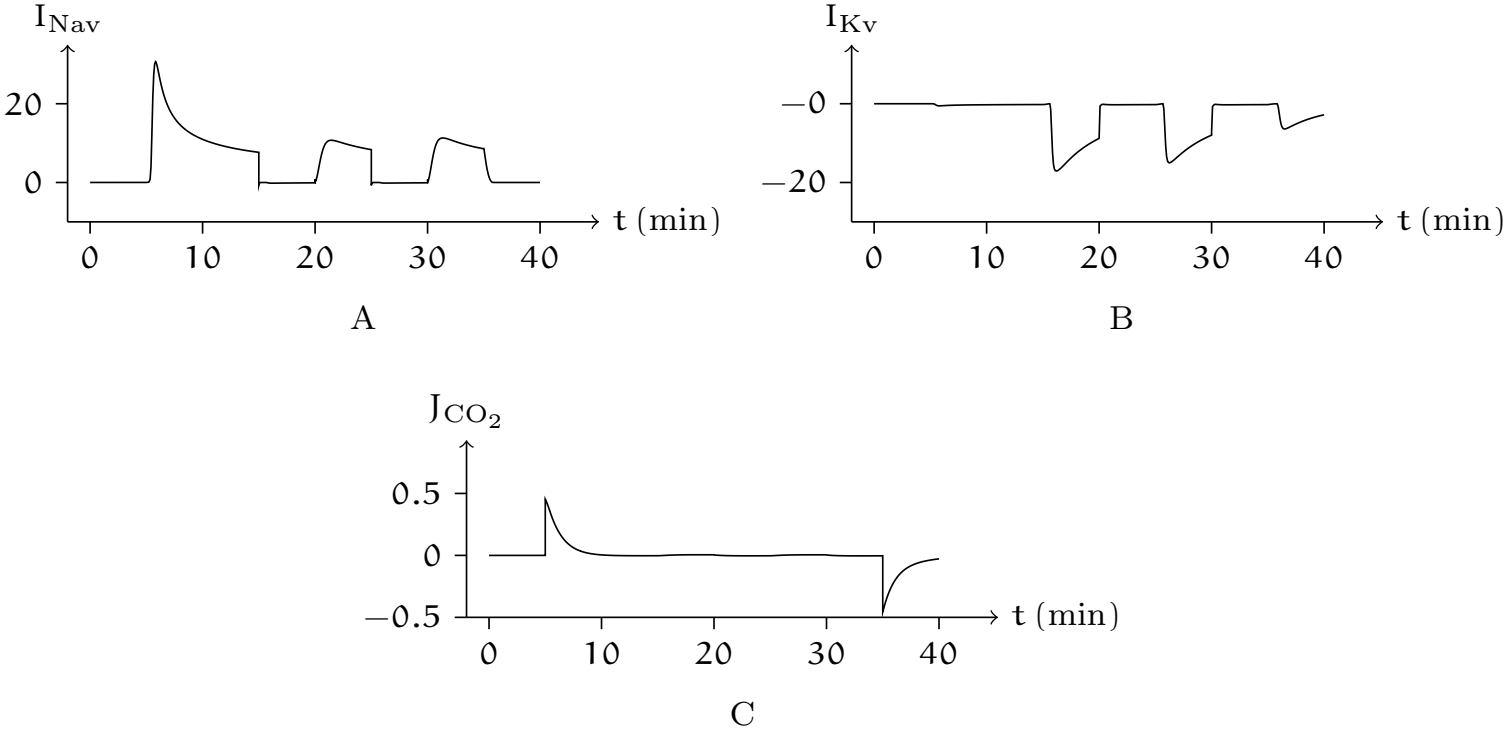
Numerical simulation of ionic currents in the physiological experiment involving heterologous expression of rkNBC in *Xenopus* oocytes. The data in this figure and in panels A and B of Figure 10 were obtained from the same numerical simulation. (A) Time course of voltage-gated sodium current. (B) Time course of voltage-gated potassium current. (C) Time course of carbon dioxide flux via simple diffusion.

## 6 Discussion

### 6.1 Cellular Homeostasis

The conventional mechanistic framework conceptualizes symporters as alternating-access “coupled transport machines”: substrate binding triggers conformational transitions that harness the thermodynamically favorable movement of one solute down its electrochemical gradient to drive the stoichiometric, concerted translocation of a second solute across the membrane.

Within this framework, transport kinetics are formulated predominantly as functions of electrochemical gradients and substrate concentrations. This framework explains how substrates trigger conformational changes and drive transmembrane flux, yet in its canonical form it lacks an intrinsic mechanism to sense intracellular homeostatic signals, and thus cannot describe autonomous modulation of transport activity without invoking additional regulatory layers.

In this work, we propose a homeostasis-centered complement to the conventional view: a core physiological function of symporters is to actively regulate and stabilize the intracellular concentration of specific solutes by coupling their transport to the electrochemical gradients of co-transported partners. Our simulation results demonstrate that incorporating an intracellular concentration-dependent gating mechanism suffices to recapitulate a broad spectrum of experimentally observed physiological behaviors across multiple evolutionarily distinct symporter families.

### 6.2 Activated, unactivated, and inactivated

Our model distinguishes among the activated, unactivated, and inactivated states as follows: A symporter resides in or tends toward the unactivated state if and only if it satisfies 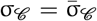. An inward-permissive (outward-permissive) symporter resides in the activated state if and only if it mediates inward (outward) cotransport; otherwise, it resides in the inactivated state.

Taking NKCC as an example, our model explains the relevant physiological experimental phenomena as follows: Fitting the physiological experimental data yields the parameter 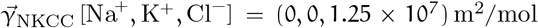. Thus, intracellular chloride deficiency places NKCC in an inward-permissive state (see Figure 7). Consequently, when extracellular concentrations of sodium, potassium, and chloride ions are sufficiently high, NKCC stably mediates inward cotransport (see Figure 8). This observation may superficially suggest that extracellular sodium and potassium activate NKCC’s inward cotransport; yet they serve solely as substrates, not as agonists, therefore eliciting no stimulus. Conversely, excess intracellular chloride ions place NKCC in an outward-permissive state. If, under this condition, extracellular sodium, potassium, and chloride ions remain excessively abundant, thereby precluding NKCC from mediating outward cotransport, NKCC enters an inactivated state. Accordingly, inhibition of NKCC transport by high intracellular chloride is observed (see Figure 7). If instead extracellular sodium, potassium, or chloride ions are severely deficient while intracellular sodium and potassium ions are abundant,

NKCC stably mediates outward cotransport (see Figure 9). Moreover, the rate of outward cotransport increases with increasing intracellular sodium and potassium ion concentrations.

### 6.3 Voltage dependence

Symporters that are sensitive to charged ions necessarily exhibit voltage dependence. This arises because the transmembrane electric field modulates the local concentration of charged ions at the cytoplasmic membrane surface. In particular, when a symporter exhibits sensitivity to intracellular cations—that is, when its cation-sensitivity coefficient exceeds zero—membrane depolarization inhibits its inward transport activity by driving cations toward the cytoplasmic membrane surface. This phenomenon has been experimentally and computationally demonstrated in both the sodium-glucose symporter (SGLT) and the sodium-phosphate symporter (NaPi) (see Figures 1 and 4). For SGLT1, quantitative fitting of electrophysiological data yields 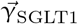[Na^+^, αMDG] = (1.5 *×* 10^9^, 0) m^2^/mol, indicating that regulation occurs via Na^+^ binding at the cytoplasmic surface, not via αMDG. Consequently, membrane depolarization elevates the local Na^+^ concentration at the cytoplasmic membrane surface, decreasing the population of inward-permissive conformational states, and ultimately reducing the net inward transport flux. Because voltage dependence in such symporters is mechanistically rooted in ion-sensitive conformational transitions, voltage-dependent symporters invariably exhibit measurable sensitivity to specific charged ions.

### 6.4 Step delineation

In general, calculating net transport rates does not necessitate highly granular step delineation. However, two specific conditions preclude step merging: (i) a large disparity in transport coefficients between consecutive steps; and (ii) the presence of a kinetic branch point within the transport cycle. In both cases, the steps must be treated as distinct and parameterized with quantitative rigor.

For instance, in SGLT1-mediated inward cotransport, Na^+^ binding precedes αMDG binding: two Na^+^ ions bind first, followed by one αMDG molecule. Given the substantial difference in binding affinities between Na^+^ and αMDG, kinetic modeling requires at least two sequential steps to ensure mechanistic fidelity. Moreover, the model parameters constrained by physiological experimental data satisfy κ[SGLT1_i1_; SGLT1_i3_(Na^+^, Na^+^)] = κ[SGLT1_e1_; SGLT1_e3_(Na^+^, Na^+^)] *≪* κ[SGLT1_i3_ (Na^+^, Na^+^); SGLT1_e3_(Na^+^, Na^+^)], hence SGLT1-mediated inward cotransport exhibits strong dependence on extracellular αMDG but negligible dependence on extracellular Na^+^ (see Figure 2).

For example, the sodium-coupled phosphate symporter accommodates bidirectional cation selectivity: it may bind Na^+^ or Li^+^ first, necessitating branching steps in the kinetic scheme. The model parameters constrained by physiological experimental data satisfy 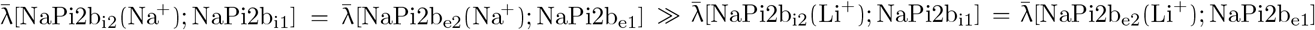, and 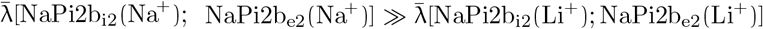, hence the pronounced inhibitory effect of Li^+^ on NaPi2b-mediated inward cotransport (see Figure 5) likely arises from Li^+^’s markedly slower dissociation rate relative to Na^+^.

### 6.5 Functional crosstalk

Transporters and ion channels functionally cooperate to maintain cellular electro-chemical homeostasis.

Ion channels typically operate on faster timescales than secondary active transporters such as symporters. Consequently, modulation of symporter activity by ion channel-mediated changes in membrane potential or intracellular ion concentrations is a well-established regulatory mechanism. Specifically, ion channel gating alters transmembrane ion gradients and membrane potential, thereby influencing the ther-modynamic driving force and conformational equilibrium of coupled transporters. In contrast, transporter activity can indirectly regulate ion channel function by driving time-dependent shifts in intracellular ion concentrations and membrane potential. The reciprocal interplay between transporters and ion channels introduces substantial mechanistic complexity, necessitating quantitative modeling to dissect their integrated physiological functions.

Using quantitative modeling constrained by electrophysiological and intracellular pH measurements in *Xenopus laevis* oocytes expressing rkNBC, we find that extracellular application of 1.5%CO_2_/10 mM 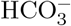 does not directly activate rkN-BC. Rather, CO_2_ crosses the plasma membrane via simple diffusion (permeability coefficient 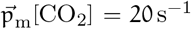; see Figure 12C), elevating intracellular 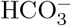, which promotes transition of rkNBC into an outward-permissive conformation (see Figure 11C). However, under the current ionic conditions, which include physiological extracellular Na^+^ and 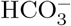, the thermodynamic constraints restrict rkNBC to net inward transport. Therefore, upon addition of CO_2_ (t > 5 min), intracellular acidification is attenuated relative to control (see Figure 10, B and D), concomitant with cellular hyperpolarization (see Figure 10A); moreover, the majority of rkNBCs rapidly transition into an inactivated statenamely, the interrupted transport state explicitly implemented in the model (see Figure 11C). Thereafter, the subsequent slow recovery of membrane potential (t < 15 min) is mediated exclusively by sodium influx through voltage-gated sodium channels (see Figure 12A).

At t = 15 min, abrupt removal of extracellular sodium ions induces immediate inactivation of sodium channels and concurrent reactivation of the rkNBC symporter from its inactivated state, thereby enabling outward cotransport; this leads to a decrease in intracellular pH and a pronounced depolarization that activates voltage-gated K^+^ channels, eliciting an outward K^+^ current that counterbalances the inward current mediated by rkNBC (see Figure 12B). Consequently, membrane potential stabilizes over the interval t/min *∈* [15, 20].

These interdependent, time-resolved dynamics underscore that transporter–channel crosstalk cannot be unambiguously elucidated by intuitive analysis of experimental recordings alone; such crosstalk emerges rigorously only when physiological behavior is interpreted through a mechanistic, multi-parameter computational framework.

## 7 Conclusion

This paper presents a general mathematical framework for modeling symport. The development of such a framework requires addressing two fundamental challenges:

First, abstracting and characterizing the shared dynamical essence common to diverse symporter families. Although symporters differ significantly in sequence and conformation, general modeling requires that they all satisfy the same functional paradigm. Therefore, the general model must abstract away nonessential structural details and establish a broadly applicable dynamical framework that satisfies thermo-dynamic conservation laws and respects the topology of conformational transitions.

Second, achieving a quantitative cross-scale mapping from single-molecule conformational dynamics to cellular physiological functions. At the microscopic level, symporters execute substrate recognition, binding, translocation, and release through conformational transitions; at the macroscopic level, their function manifests as cellular uptake of various solutes, maintenance of pH and ion homeostasis, regulation of osmotic balance, modulation of membrane potential, and control of metabolic states. Establishing a rigorous analytical link between these levels requires assigning well-defined parameters to key conformational states and ensuring that the conformational distribution quantitatively reproduces experimentally measurable physiological outputs—including transport rates, steady-state currents, pH profiles, and inhibition kinetics—which constitutes the core task of scale bridging.

This article is grounded in the fundamental physiological principle of cellular homeostasis and proposes a core hypothesis for symporter gating: transport proteins regulate intracellular concentrations of specific solutes within thermodynamically feasible bounds. Based on this hypothesis, we formulate a general mathematical model of symporters, built upon minimal conformational cycles as its topological scaffold and incorporating probabilistic transition rates to quantitatively describe conformational interconversion kinetics.

We rigorously validated this model by simulating published experimental data from prototypical members of the SGLT, NaPi-II, NKCC, and NBC transporter families. Results demonstrate that a single unified parameter set achieves robust fits across diverse experimental conditions—including varying membrane potentials and substrate concentration gradients. Moreover, integration of this model with our previously established general ion channel model establishes a quantitative framework for both simulating transporter–channel crosstalk and dissecting its underlying functional mechanisms.

Our next objective is to generalize this approach to uniporters and antiporters, thereby establishing a unified mathematical framework for all secondary active transporters—one that rigorously reconciles mechanistic interpretability with quantitative predictive capability.

## Footnotes

1 In the past, studies of membrane potential often overlooked the interaction between ion distribution and the membrane electric field. Early models, such as the Hodgkin-Huxley model [1], emphasized ion conductances but assumed uniform intracellular/extracellular ion concentrations. This simplification neglected spatial ion gradients and dynamic concentration changes during prolonged cellular activity. To address this limitation, researchers have adopted computational modeling approaches, including finite-element simulations, which now integrate three-dimensional ion gradients and subcellular compartmentalization.

2 4-deoxy-4-fluoro-D-glucopyranoside.

3 For example, in NKCC2, intracellular chloride depletion is sensed not by the transporter itself but by an upstream kinase cascade, in which WNK3 lies upstream of SPAK and has been proposed as the chloride-sensitive kinase [18]. The signal is relayed through a WNK3–SPAK complex and results in phosphorylation of conserved N-terminal threonines (T96, T101, and T111); alanine substitution of all three residues or expression of catalytically inactive WNK3 prevents activation by low [Cl^−^]_i_, whereas disruption of the WNK3–SPAK interaction abolishes WNK3-induced activation.

4 As a representative example, the stoichiometric cotransport ratio mediated by NKCC is 1 Na^+^ : 1 K^+^ : 2 Cl^−^.

5 For instance, in electrophysiological recordings that use continuous perfusion systems, such as patch-clamp experiments, or in microfluidic chip platforms, the extracellular environment is typically maintained and exchanged via precisely regulated fluid delivery mechanisms.

6 SGLT2 is expressed poorly in heterologous expression systems, where the rate of transport is generally orders of magnitude less than that for SGLT1. Human SGLT3 is not a transporter but instead a glucose sensor.

7 ∥SGLT1∥ = 10 nmol/m^2^, α_SGLT1_ = 200 s^−1^, ν_−A SGLT1_ = 20 s^−1^, ν_−B SGLT1_ = 20 s^−1^, 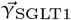 [Na^+^, αMDG] = (1 *×* 10^8^, 0) m^2^/mol.

8 The quantity 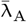 has units of s^−1^, and κ has units of mol m^−2^. These units are defined herein and will not be repeated in subsequent equations or text.

9 ∥NaPi2b∥ = 0.1 nmol/m^2^, α_NaPi2b_ = 10 s^−1^, ν_−A NaPi2b_ = 1 s^−1^, ν_−B NaPi2b_ = 1 s^−1^, 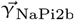 [Na^+^, Li^+^, P_i_] = (1.5 *×* 10^9^, 0, 0) m^2^/mol.

10 ∥NKCC∥ = 10 nmol/m^2^, α_NKCC_ = 1 s^−1^, ν_−A NKCC_ = 1 s^−1^, 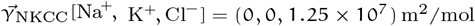.

11 α_rkNBC_ = 10 s^−1^, ν_−A rkNBC_ = 1 s^−1^, ν_−B rkNBC_ = 1 s^−1^, 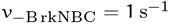 [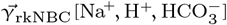] = (0, 0, 1*×* 10^12^) m^2^/mol.

## Notes

### Competing Interest Statement

The authors have declared no competing interest.

### Summary of Updates

The title, analysis, and discussion sections have been revised to more accurately represent the scope, methodology, and conceptual contributions of our work.

